# A regulatory gene network that couples floral transition to shoot apical meristem morphology in Arabidopsis

**DOI:** 10.1101/2024.05.10.593152

**Authors:** Enric Bertran Garcia de Olalla, Gabriel Rodríguez-Maroto, Martina Cerise, Alice Vayssières, Edouard Severing, Yaiza López Sampere, Kang Wang, Sabine Schäfer, Pau Formosa-Jordan, George Coupland

## Abstract

Plants flower in response to environmental signals. These signals change the shape and developmental identity of the shoot apical meristem (SAM), causing it to form flowers and inflorescences rather than leaves. How the changes in SAM shape and identity are coordinated is poorly understood. Using genetics, confocal microscopy and RNA sequencing, we show that reciprocal repression of the APETALA2 (AP2) and SUPPRESSOR OF OVEREXPRESSION OF CONSTANS 1 (SOC1) transcription factors in the SAM is crucial in coupling these processes. During vegetative development, AP2 represses *SOC1* transcription to delay floral transition. Later, *SOC1* is activated during floral transition in response to environmental cues and represses *AP2* transcription. However, during floral transition and prior to its stable repression by SOC1, AP2 rapidly promotes an increase in the size, and changes in the morphology, of the SAM. Thus, direct reciprocal repression of AP2 and SOC1 couples floral transition and changes in SAM morphology, which initiate inflorescence development.

## Introduction

The shoot apical meristem (SAM) contains a population of stem cells that gives rise to all above-ground tissues. As cells are displaced from the stem cell niche, they differentiate and form organs on the flanks of the SAM, but the stem cell population is maintained throughout the life of the plant to allow continuous organ production^1,2^. The structure and function of the SAM change with plant age and in response to environmental signals^3,4^. Notably, during the transition to flowering, the SAM enlarges and changes identity to initiate flowers instead of leaves^5–7^. The increase in SAM size during floral transition persists in the inflorescence meristem, and likely contributes to the number of flowers formed^5,8–10^. However, how changes in meristem shape and identity are temporally coordinated remains unclear.

Exposure of Arabidopsis plants to long days (LDs) causes the SAM to transition rapidly from the vegetative to the reproductive state, which involves a radical reprogramming of the SAM transcriptome^11,12^. During this process, genes encoding transcription factors that repress flowering are downregulated, whereas the expression of floral promoters is upregulated. One of the earliest induced genes is *SUPPRESSOR OF OVEREXPRESSION OF CONSTANS 1* (*SOC1*), which encodes a MADS-domain transcription factor and promotes floral transition at the SAM^13–15^. During floral transition, the SAM also increases in size and changes in shape^5,16–19^. These changes involve faster growth in the vertical direction, increasing SAM height, than in the lateral direction, increasing width, and this imbalance creates a characteristic dome-shaped SAM. This domed shape is fully acquired before floral primordia are formed, and results in a mature inflorescence SAM that is larger than the vegetative SAM^5,16–19^. Mutations in *SOC1* delay doming of the SAM^5^, in addition to delaying flowering.

Although several mechanisms that influence SAM size have been described^20–29^, those responsible for altering SAM morphology during floral transition remain to be elucidated. The APETALA 2 (AP2) transcription factor, which confers sepal and petal identity^30,31^, is expressed in the SAM, and *ap2* mutants are early flowering and have smaller meristems at the embryonic and inflorescence stages^8,21,32^. By contrast, gain-of-function *AP2* transgenes that are resistant to microRNA172 (miR172) confer an increase in inflorescence meristem size and late flowering^8,21^. A feedback loop between the WUSCHEL (WUS) homeodomain transcription factor, expressed in the organizing centre below the stem cell niche, and the CLAVATA3 (CLV3) peptide, expressed in the central zone, maintains the size of the SAM^20,33–36^. AP2 can increase *WUS* transcription^21,37–39^ and its ectopic expression in the CLV3 or WUS domains increases SAM size^8^.

Here, we show that in Arabidopsis, AP2 and SOC1 transcription factors mutually repress each other’s expression in the SAM and that this regulatory motif ensures that AP2 strongly increases SAM size and alters its morphology specifically during floral transition.

## Results

### AP2 positively regulates meristem area during floral transition

AP2 increases the size of the inflorescence SAM and delays flowering; therefore, its effects on SAM area and shape during floral transition were examined. No significant difference between the SAM area of *ap2-12* mutants and Col-0 was observed after vegetative growth for 2 weeks under non-inductive short days (2 wSD; **Fig. 1**a–b). Plants were then transferred to long-day (LD) conditions to induce floral transition, and SAM size was measured at regular intervals until 11 LDs. The area of the Col-0 SAM increased by 5.7 fold during exposure to LDs (i.e. +11 LD vs. 2 wSD), but reached its maximum at +7 LD (9.0-fold larger). The SAM area of *ap2-12* mutants was less than half that of Col-0 at +7 LD, and therefore increased to a much lesser extent than that of Col-0 during floral transition. These results indicate that AP2 does not detectably affect SAM area during vegetative growth under SDs, but contributes to the rapid increase in SAM area observed during floral transition after exposure to LDs.

**Fig. 1.**
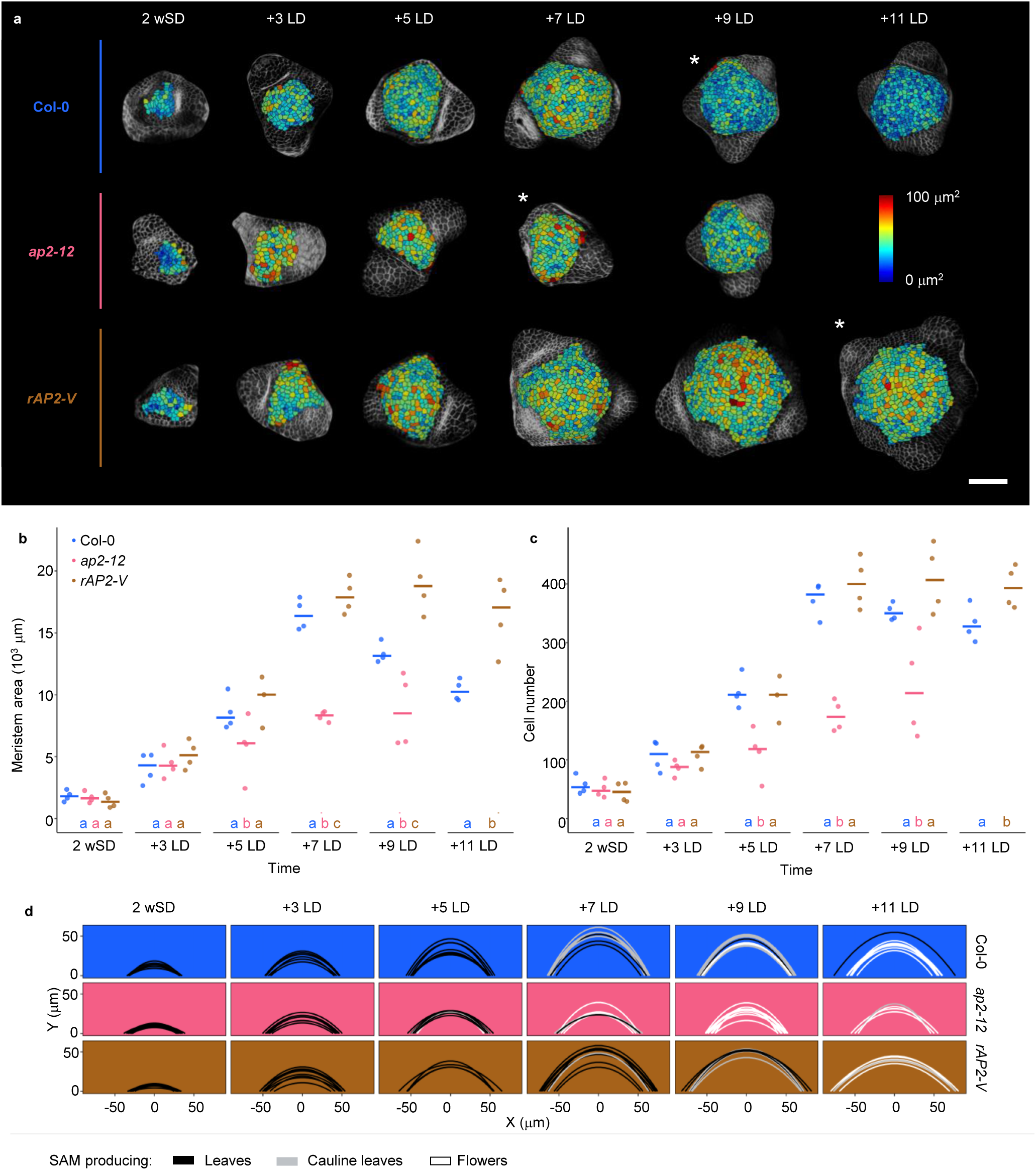
AP2 is a positive regulator of SAM size during floral transition. (a–c) Segmentation analyses of Col-0, *ap2-12* and *rAP2-VENUS* SAMs from 2-week-old plants grown under short day (SD) conditions (2 wSD) and then transferred to continuous long days (LDs). (a) Top view of the heatmap quantification of cell area in the meristem region. White asterisks indicate the first time point at which floral primordia were detected in the analysis of the corresponding genotype. Scale bar = 50 μm. (b–c) Quantification of (b) meristem area and (c) cell number. The horizontal bars represent the median value for each genotype. Significant differences among genotypes within each time point were determined via one-way ANOVA, followed by Tukey post-hoc comparisons (*p* < 0.05). Data sets that share a common letter do not differ significantly. The colour of the dots and the letters correspond to the genotype. *N* = 4 (except *rAP2-VENUS* at +5 LD, *N* = 3). (d) SAM morphology adjusted to parabolas. The parabolas are coloured according to the identity of primordia that were formed at the SAM periphery. *N* = 4– 15.

AP2 is negatively regulated by miR172 in the inflorescence SAM^40^. To test whether persistent AP2 levels above those of Col-0 further increases SAM size during floral transition, apices from plants carrying a miR172-resistant version of *AP2* tagged with VENUS and expressed from the *AP2* promoter (*rAP2-V*), were analysed (**Fig. 1**a-b). These plants exhibited enlarged inflorescence meristems compared with Col-0^8^. The SAM area of *rAP2-V* plants was comparable to that of Col-0 at 2 wSD, and the SAM area of *rAP2-V* plants increased at a comparable rate to that of Col-0 after exposure to LDs (**Fig. 1**b). However, the area of the *rAP2-V* SAM did not decrease soon after reaching a maximum size at +7 LD, as observed for Col-0^5^, but remained approximately constant at +9 and +11 LD. Therefore, the area of the *rAP2-V* SAM was 1.6-fold larger than that of Col-0 at +11 LD. These results support a transient positive role of AP2 in increasing the SAM area of Col-0 plants during floral transition, and suggest that extending the duration of its expression leads to a larger SAM.

In Col-0, enlargement of the SAM during floral transition is associated with increases in both cell number and cell area in the epidermis, hereafter referred to as Layer 1 (L1), which reach their maximum at +7 LD^5^. Therefore, these parameters were measured in *ap2-12* and *rAP2-V* plants (**Fig. 1**a, c, Supplementary Figure 1a). The SAM L1 of *ap2-12* mutants contained roughly half (47%) the number of cells of that of Col-0 at +7 LD, whereas the *rAP2-V* SAM L1 contained significantly more cells at +11 LD than that of Col-0 (Fig. 1a, c). Moreover, at the same time points, cell area was significantly increased in *ap2-12* and *rAP2-V* compared with Col-0 (Supplementary Figure 1a). Therefore, AP2 regulates both cell number and cell size in the L1 during floral transition.

### AP2 increases SAM height and width during floral transition

The height and width of *ap2-12* and Col-0 SAMs were measured to quantify differences in SAM morphology (Supplementary Figure 2a–g) and these measurements were used to reconstruct SAM shape as a parabola^41,42^ (**Fig. 1**d). Both height and width of the Col-0 SAM increased during the time course (Supplementary Figure 2c,f–g), but mean SAM width increased 1.7 fold from 2 wSD to +7 LD, whereas SAM height increased on average by almost 3.9 fold (Supplementary Figure 2c). Compared with Col-0, the SAMs of *ap2-12* mutants were significantly reduced in height from +5 LD onwards and in width from +7 LD (Supplementary Figure 2f–g). At +7 LD the *ap2-12* SAM showed a reduction of 47.8% in height and did not show the characteristic domed shape of the Col-0 SAM (**Fig. 1**d, Supplementary Figure 2g).

By contrast, the width of *rAP2-V* SAMs was greater than that of Col-0 SAMs at the two time points (i.e. +9 and + 11 LD) at which *rAP2-V* meristems were increased in size (**Fig. 1**a–b, Supplementary Figure 2f), but the height of *rAP2-V* and Col-0 meristems were similar throughout floral transition. These data demonstrate that AP2 is required in Col-0 to increase SAM width and height to confer a fully domed SAM during floral transition, and suggest that persistence of AP2 at late stages of the floral transition due to insensitivity to miR172 increases SAM width after transfer from SDs to LDs but does not further increase SAM height.

In addition to its effects on SAM development, *AP2* is a negative regulator of flowering time^32^. The early flowering of *ap2-12* mutants and the late flowering of *rAP2-V* plants were detected microscopically by scoring the time point at which floral primordia were first visible (**Fig. 1**a, d; Supplementary Figure 1b–d, Supplementary Figure 2h–j). In Col-0 this occurred at +9 LD, 2 days after the SAM reached maximum height. Furthermore, cauline leaf primordia of Col-0 were first detected based on their morphology at 7 LDs when SAM height was at its maximum (**Fig. 2**d, Supplementary Figure 2g). The SAM of *rAP2-V* plants reached maximum height at the same time point as Col-0 (+7 LD), and cauline leaf primordia were also first detected at that stage (**Fig. 1**d, Supplementary Figure 2a, g). However, floral primordia were first detected later in *rAP2-V* plants, at +11 LD, and although the height of *rAP2-V* SAM reduced after its peak at +7 LD in a similar way to that of Col-0, the *rAP2-V* SAM increased in width until +11 LD (Fig. 1d, Supplementary Figure 2f, g).The majority of *ap2-12* mutants (>50%) already exhibited floral primordia at +7 LD, although they did not show the large increase in SAM height observed for the SAM of Col-0 at the same stage. These observations demonstrate a temporal sequence of events in Col-0 in which meristem size increases and then decreases prior to floral development, and that in *ap2-12* mutants floral development occurs independently of these changes in SAM size.

**Fig. 2.**
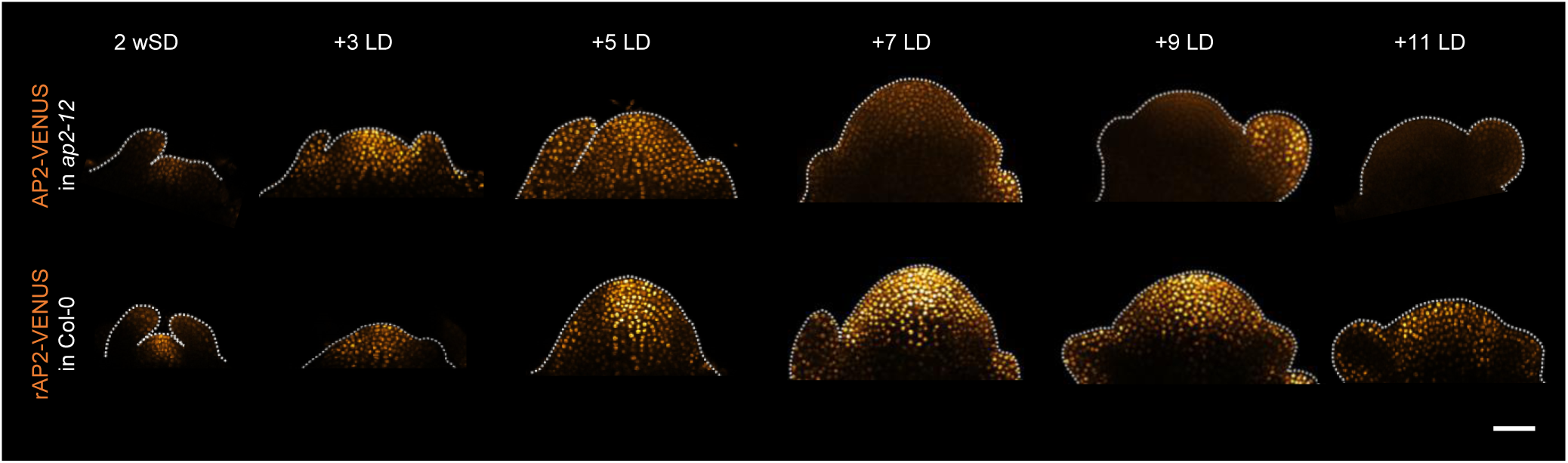
AP2 protein is present in the SAM during floral transition. The pattern of protein accumulation of *AP2::AP2:VENUS* in *ap2-12* and *AP2::rAP2:VENUS* in Col-0 at the SAM of 2-week-old plants grown under short day (SD) conditions (2 wSD) and then transferred to continuous long days (LDs). The shape of the acquired meristem and its peripheral organs is indicated with a dotted white line. Scale bar = 50 μm.

### AP2 protein is present in the SAM during floral transition

AP2:VENUS (AP2-V) fusion protein expressed from the *AP2::AP2:VENUS* transgene is present in the SAM during vegetative development and absent from the inflorescence SAM^8,40^, but its expression during floral transition has not been studied. Confocal microscopy was used to image AP2-V in the SAM of *ap2-12 AP2::AP2-VENUS* plants grown for 2 wSD and then transferred to LDs. AP2-V was detectable from 2 wSD to +7 LD, but at +9 LD when AP2-V was clearly detected in floral primordia, its abundance in the SAM was very low (Fig. 2, Supplementary Figure 3). Therefore, although AP2 delays floral transition during vegetative development, it is still present in the SAM during floral transition at +5 to +7 LD when it is required for the SAM to reach its maximum height. Moreover, rAP2-V was detectable in the SAM until the end of the time course at +11 LD, and therefore its extended presence in the SAM at +9 and +11 LD correlates with the time points at which the SAM of rAP2 plants is increased in width and size compared with that of Col-0 plants. Overall, these data, together with the phenotypic analysis of the SAMs of the different genotypes, support the hypothesis that AP2 acts directly in the SAM during floral transition to increase SAM size and alter its morphology.

### AP2 delays floral transition by repressing *SOC1* expression before floral transition

To understand how AP2 regulates SAM height and width during floral transition, global gene expression analysis was performed by RNA sequencing (RNA-Seq) using apices of Col-0 and *ap2-12* mutants grown under continuous LD. First, the morphological analyses of the SAMs of *ap2-12*, Col-0 and *rAP2-V* were repeated under continuous LDs to test whether the SAMs developed similarly to those of plants transferred from SD to LD. Col-0 formed a SAM with maximum height at 14 LD (Fig 3a; Supplementary Figure 4a) and *ap2-12* exhibited a smaller meristem from 14 LD onwards (Supplementary Figure 5a–b), whereas the meristem of *rAP2-V* became enlarged from 12 LD. The smaller meristem of *ap2-12* mutants was again associated with a reduction in cell number (Supplementary Figure 5c). At 14 LD, when the Col-0 SAM showed maximum height, the *ap2-12* SAM exhibited a reduction in height and width of 50% and 25%, respectively, with respect to Col-0 (Supplementary Figure 4). Moreover, although the height of *rAP2-V* meristems was comparable to that of Col-0 at 14 LD, their width was greater and reached a maximum at 17 LD. Overall, the SAMs of Col-0, *ap2-12* and r*AP2-V* plants showed similar differences in morphology under continuous LDs to those described after transfer from 2w SD to LD.

**Fig. 3.**
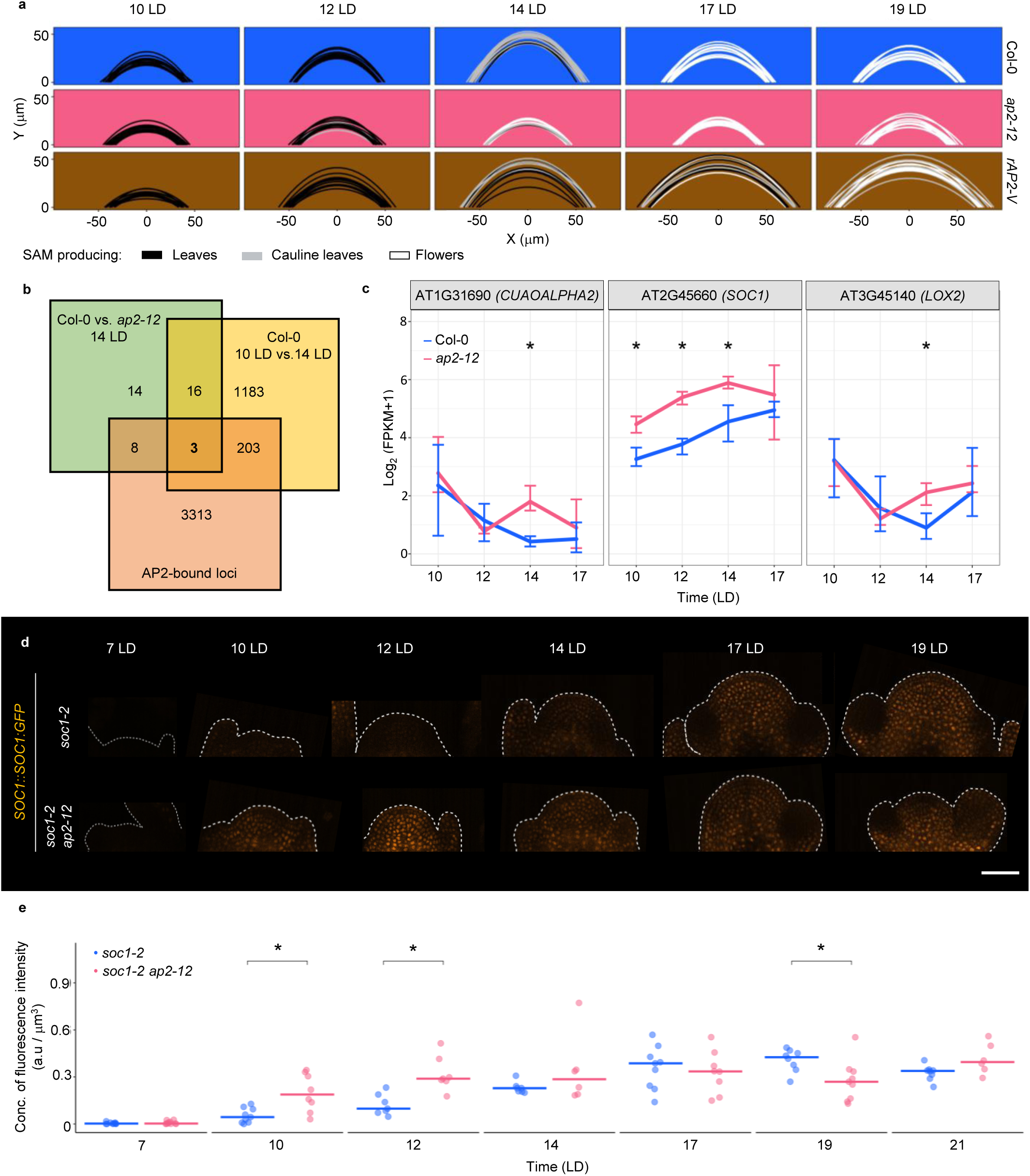
AP2 is a negative regulator of *SOC1* expression at the SAM before and during floral transition. (a) SAM morphology of Col-0, *ap2-12* and *rAP2-VENUS.* The parabolas are coloured according to the identity of primordia that were formed at the SAM periphery. *N* = 10–21. (b–c) Global transcriptome profiling via RNA-Seq of dissected meristems of Col-0 and *ap2-12* plants. (b) Venn diagram showing the overlap between the list of differentially expressed genes (DEGs) at 14 long days (LDs) between Col-0 and *ap2-12*, the list of DEGs in Col-0 between 10 LD and 14 LD, and the AP2-bound loci^32^. (c) Expression profiles under LDs for dissected plant apices of the genes that are present in the three lists compared in b. The gene symbol is represented above each plot. Error bars represent the range between the maximum and minimum values among the three replicates. Significant differences between genotypes at the same time point were determined via likelihood ratio test. Statistical differences between Col-0 and the other genotypes are indicated with an asterisk (adjusted *p-*value < 0.05). (d) Pattern of protein accumulation of *SOC1::SOC1:GFP* at the SAM in *soc1-2* and *soc1-2 ap2-12* plants under continuous LDs. The shape of the acquired meristem and its peripheral organs is indicated with a dotted white line. Scale bar = 50 μm. (e) Quantification of SOC1:GFP concentration of fluorescence intensity (total fluorescence divided by volume) at the shoot apex (from the tip to 50 μm deep in the basal direction) in *soc1-2* and *soc1-2 ap2-12* mutant backgrounds during continuous LDs. The dots are coloured according to the mutant background of the analysed plant. The horizontal bars represent the median value for each genotype. Comparisons within each time point between genotypes were performed via Wilcoxon rank sum test. Statistical significance (*p* < 0.075) is indicated with an asterisk. *N* = 6–10. The quantification of concentration of fluorescence intensity is consistent when performing the analysis with a depth of 20 μm, as shown in Supplementary Figure 6b.

Apices of Col-0 and *ap2-12* plants grown under continuous LDs for 10, 12, 14 and 17 LD were then harvested for RNA-Seq analysis. This approach yielded 103 genes that were differentially expressed (DEGs) in *ap2-12* compared with Col-0 at one or more time point (Supplementary Table 1). To identify candidate genes for regulators of meristem morphology, DEGs between SAMs that had reached their maximum height and those that had not were selected by comparing the RNA-Seq data between genotypes and time points. Specifically, the smaller SAM of *ap2-12* mutants at 14 LD was compared with the SAM of Col-0 that had increased in height and width at the same time point (Supplementary Table 2), and these DEGs were cross-referenced with those identified by comparing the SAM of Col-0 at 14 LD with the SAM of Col-0 at 10 LD before it increased in height and width (Supplementary Table 3). The resulting list of DEGs common to both comparisons (Supplementary Table 4) was then compared with the list of direct target genes of AP2 identified by ChIP-Seq analysis^32^ (Supplementary Table 5). This approach identified only three candidate genes (**Fig. 3**b): *LIPOXYGENASE 2, COPPER AMINE OXIDASE ALPHA 2* and *SOC1.* In subsequent analyses, we focused on *SOC1*, because it is a positive regulator of flowering time that acts downstream of the main effector of the photoperiodic flowering pathway, *FLOWERING LOCUS T*^13,43^, and SOC1 protein accumulates at the SAM during floral transition and persists in the inflorescence meristem^44^. Moreover, in the RNA-Seq data, *SOC1* is one of the earliest responsive genes to the loss-of-function of *AP2,* exhibiting a higher level of expression already at 10 LD in *ap2-12* mutants compared with Col-0 (**Fig. 3**c).

To compare the temporal and spatial patterns of SOC1 protein accumulation at the SAM in *ap2-12* and Col-0 SAMs, the *SOC1::SOC1:GFP* reporter was introduced into *soc1-2* and *soc1-2 ap2-12* mutant backgrounds (**Fig. 3**d–e; Supplementary Figure 6). At 10 LD and 12 LD, SOC1:GFP was detected more strongly at the SAM in the *ap2-12* mutant background (*ap2-12 soc1-2 SOC1::SOC1:GFP*) than in wild-type plants (*soc1-2 SOC1::SOC1:GFP*), indicating that AP2 represses *SOC1* transcription in the wild-type SAM during vegetative development. However, by 14 LD SOC1 abundance had increased in the SAM of wild-type plants and was detected at a similar level in the SAM of *ap2-12* mutants (Fig. 3d-e). Collectively, these results indicate that AP2 negatively regulates *SOC1* expression at the SAM of wild-type plants before floral transition.

The genetic interaction between *soc1-2* and *ap2-12* in the control of flowering time was then characterized by comparing the days to bolting and the number of rosette and cauline leaves formed by *soc1-2*, *ap2-12* and *soc1-2 ap2-12* (Fig. 4a-d), thereby extending previous data for total leaf number^32^. The *soc1-2 ap2-12* double mutants formed on average 6.2 more rosette leaves than *ap2-12* and Col-0, but 5.2 fewer than *soc1-2*, and bolted 2 days later than Col-0 at a similar time to *soc1-2* (Fig. 4a-c). Collectively, these data indicate that SOC1 is an important promoter of flowering time downstream of AP2, but that other genes must also contribute because *soc1-2* is not completely epistatic to *ap2-12*. The inflorescence of *ap2-12* formed fewer cauline leaves than Col-0 and *soc1-2* formed more than Col-0, whereas *ap2-12 soc1-2* formed a similar number to Col-0 (**Fig. 4**a–c). Overall, these data demonstrate that *SOC1* is an important promoter of flowering time downstream of *AP2,* consistent with its increased abundance in the SAM of *ap2-12* mutants, whereas AP2 is important for the formation of cauline leaves in *soc1-2* and Col-0.

**Fig. 4.**
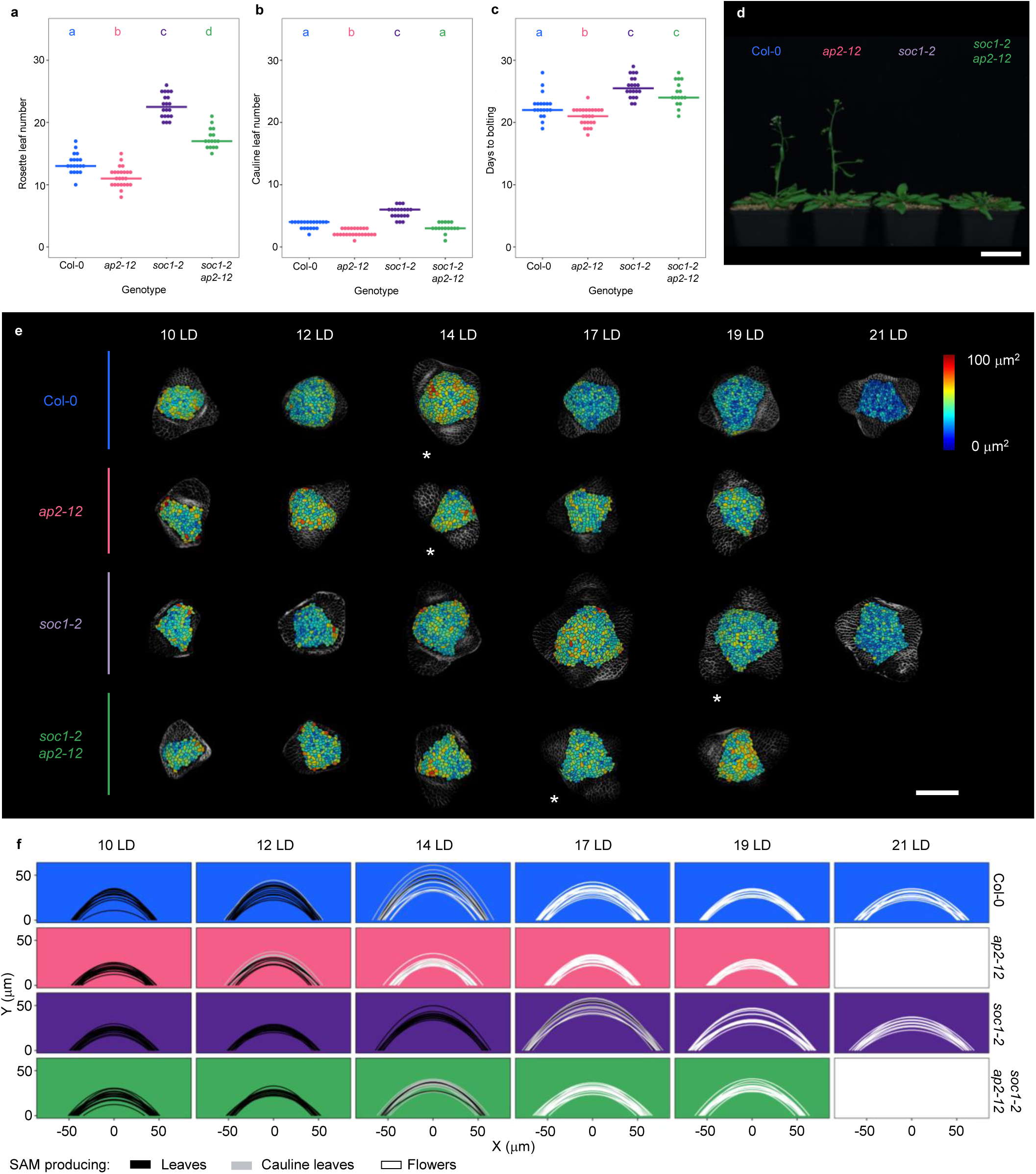
*SOC1* is an important positive regulator of flowering time downstream of *AP2.* (a–d) Flowering-time analysis of Col-0, *ap2-12*, *soc1-2* and *soc1-2 ap2-12* under long days (LDs). (a–c) Flowering time according to (a) rosette leaf number, (b) cauline leaf number and (c) days to bolting. The horizontal bars represent the median value for each genotype. Significant differences among genotypes were determined via one-way ANOVA, followed by Tukey post-hoc comparisons (*p* < 0.05). Data sets that share a common letter do not differ significantly. *N*= 16–24. (d) Photograph of representative plants of the used genotypes grown for 27 days under LD conditions. Scale bar = 5 cm. (e) SAM morphology of Col-0, *ap2-12*, *soc1-2* and *soc1-2 ap2-12* under continuous LDs. The parabolas are coloured according to the identity of primordia that were formed at the SAM periphery. *N* = 11–18. (f) Top view of the heatmap quantification of cell area in the meristem region via segmentation in SAMs of Col-0, *ap2-12*, *soc1-2* and *soc1-2 ap2-12.* White asterisks indicate the first time point at which floral primordia were detected in the analysis of the corresponding genotype. Scale bar = 50 μm. *N* = 4.

### The interaction between SOC1 and AP2 couples changes in SAM morphology to the floral transition

The increase in SAM area that occurs during floral transition is delayed in *soc1* mutants^5^. To test whether *soc1-2* meristems exhibit a delayed change in SAM morphology, the height and width of the SAM of *soc1* mutants were quantified through a time course of continuous LDs. The maximum values of SAM area, height and width for *soc1-2* meristems were detected at 17 LD, 3 days later than in Col-0 (**Fig. 4**e, Supplementary Figure 7). These results indicate that *SOC1* regulates the timing of the increases in SAM height and width that occur in Col-0 during floral transition.

We then analysed SAM morphology of *soc1-2 ap2-12* double mutants to ask whether the increase in SAM height and width that occurs later in *soc1-2* mutants is still mediated by AP2. Throughout floral transition under continuous LD, the meristem area of *soc1-2 ap2-12* was comparable to that of *ap2-12* (**Fig. 4**e, Supplementary Figure 8a). Moreover, the *soc1-2 ap2-12* double mutant showed reduced meristem height in comparison with Col-0 until 17 LD, similar height to Col-0 at 19 LD and 21 LD, and similar width to Col-0 until 21 LD (Fig. 4f, Supplementary Figure 7a, b). Thus, the increase in width and height that occurs later than in Col-0 in *soc1-2* mutants largely depends on AP2 activity.

SOC1 is a direct repressor of *AP2*^44,45^ (Supplementary Figure 9b)^45,46^, suggesting that SOC1 might be partly responsible for the lower level of AP2 accumulation in the SAM during floral transition (**Fig. 2**), and that the later increase in SAM size in *soc1-2* mutants might be associated with an extended persistence of AP2 in the SAM. To test the effect of SOC1 on AP2 expression in the SAM, *AP2::AP2:VENUS* expression was compared in *ap2-12* and *soc1-2 ap2-12* plants grown under LDs (**Fig. 5**, Supplementary Figure 10). In the *ap2-12 AP2::AP2:VENUS* background, the level of AP2:VENUS slowly decreased from 7 LD, but was still present at 14 LD when the SAM of Col-0 reaches maximum height. Thereafter, the fluorescence of AP2-V decreased more steeply in the SAM and reached its lowest levels from 17 LD to 19 LD onwards. In the *soc1-2 ap2-12 AP2::AP2:VENUS* mutant background, the temporal pattern of AP2:VENUS expression was similar to that in the *ap2-12 AP2::AP2:VENUS* background, but AP2:VENUS was present at higher levels in the SAM from 17 LD to 21 LD. Therefore, AP2 protein accumulates for longer in *soc1-2* and increases in abundance at 17 LD prior to when the *soc1-2* SAM first becomes taller than that of Col-0 at 19 LD (Supplementary Figure 7). Taken together, these experiments indicate that the repression of *AP2* by SOC1 in Col-0 restricts the increase in SAM size to the early stages of floral transition, until approximately 14 LD, and that the longer duration of *AP2* expression in *soc1-2* enables the later increase in SAM size observed in the mutant.

**Fig. 5.**
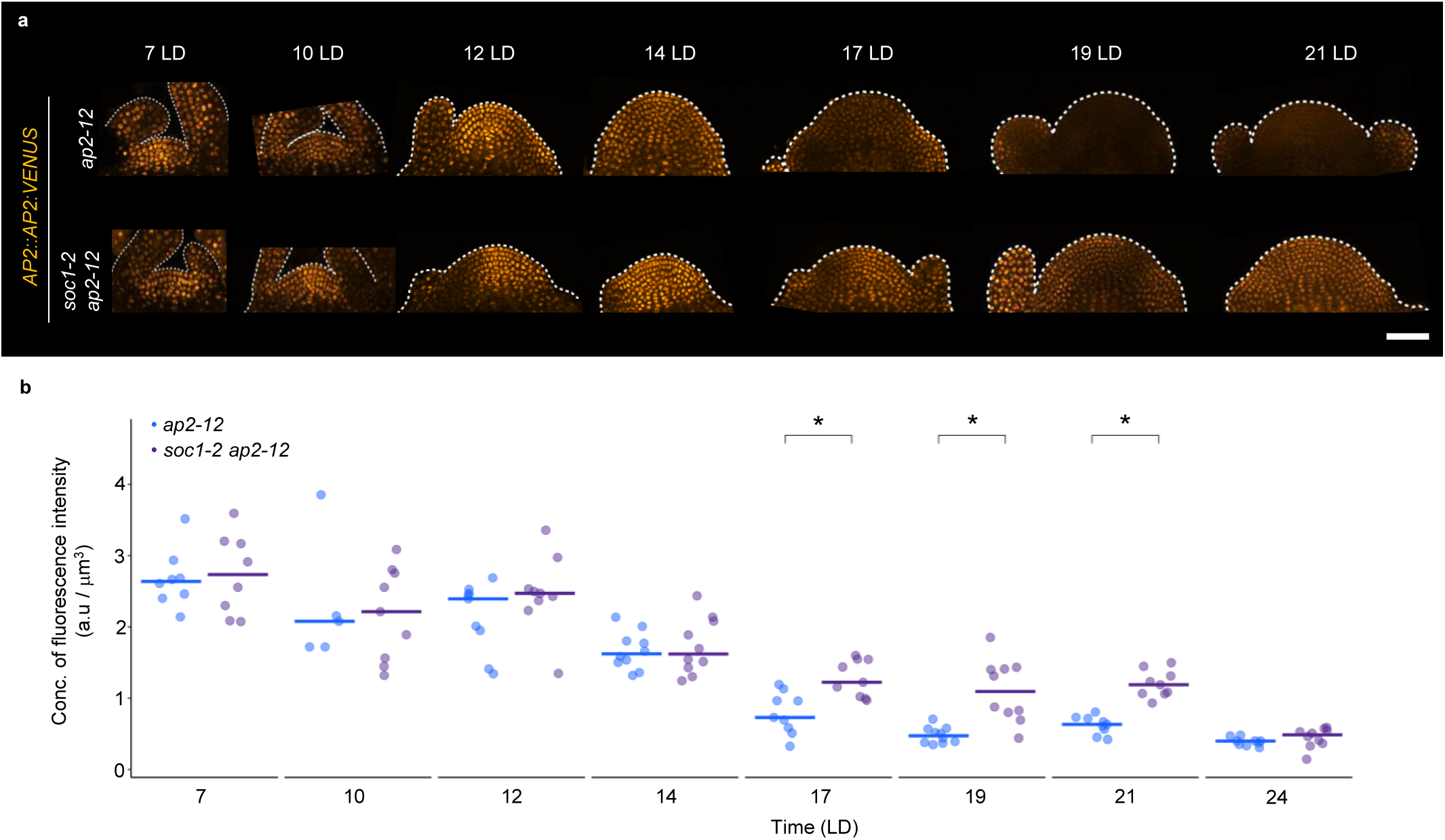
SOC1 is a negative regulator of *AP2* expression at the SAM during floral transition. (a) Pattern of protein accumulation of *AP2::AP2:VENUS* at the SAM in *ap2-12* and *soc1-2 ap2-12* plants under continuous longs days (LDs). The shape of the acquired meristem and its peripheral organs is indicated with a dotted white line. Scale bar = 50 μm. (b) Quantification of AP2:VENUS concentration of fluorescence intensity (total fluorescence divided by volume) at the shoot apex (from the tip to 50 μm deep in the basal direction) in *ap2-12* and *soc1-2 ap2-12* mutant backgrounds during continuous LDs. The dots are coloured according to the mutant background of the analysed plant. The horizontal bars represent the median value for each genotype. Comparisons intra-time point between genotypes were performed via Wilcoxon rank sum test. Statistical significance (*p* < 0.075) is indicated with an asterisk. *N* = 5–10. The quantification of concentration of fluorescence intensity is consistent when performing the analysis with a depth of 20 μm, as shown in Supplementary Figure 10b.

## Discussion

We show that reciprocal repression of expression of the AP2 and SOC1 transcription factors plays a central role in integrating morphological changes of the SAM with the acquisition of floral primordia identity. These proteins likely directly repress each other’s transcription, because each factor binds directly to the promoter of the other gene^32,44,45^. During vegetative development, AP2 delays floral transition^32^, in part by repressing *SOC1* transcription, but does not detectably influence vegetative SAM size (**Fig. 6**a). However, exposure of plants to environments such as LDs overcomes the repression of *SOC1* by AP2 (**Fig. 6**b), and a rise in SOC1 abundance represses *AP2* transcription (**Fig. 6**c). AP2 protein levels are also reduced through post-transcriptional regulation mediated by miR172^46,47^, and our analysis of *rAP2-V* indicates that insensitivity to miR172 extends the duration of AP2 expression into the mature inflorescence SAM. The reduction in AP2 level by SOC1 and miR172 further accelerates the increase in *SOC1* activity and floral transition. However, in the short time interval between the initiation of floral transition and the disappearance of AP2 from the SAM, at about 12–14 LDs after germination, AP2 promotes increases in SAM size and alterations in its morphology to a more dome-like shape (**Fig. 6**b). Therefore, the doming of the SAM associated with flowering is limited to the early stages of floral transition prior to the reduction in AP2 by the actions of SOC1 and miR172. In *soc1* mutants, floral transition and the repression of *AP2* are both delayed, and the SAM becomes domed later. Thus, the mutual repression of SOC1 and AP2 determines the timing of floral transition and ensures that the SAM rapidly increases in size only during the early stages of floral transition.

**Fig. 6.**
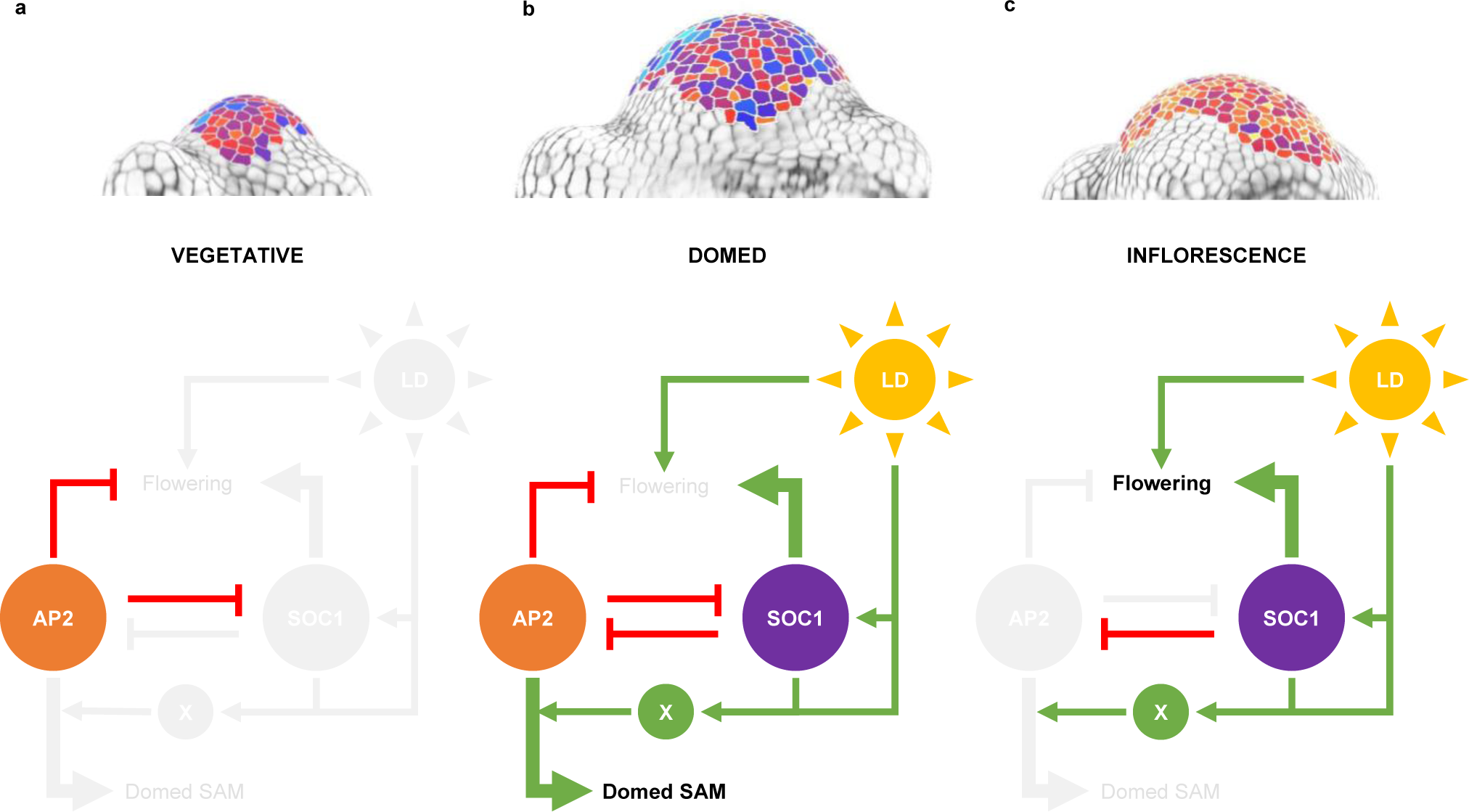
The mutual repression of *AP2* and *SOC1* couples changes in SAM morphology with floral transition. Schematic representation of AP2 and SOC1 regulation of SAM morphology and flowering at the (a) vegetative, (b) floral transition, and (c) inflorescence stages. At the vegetative stage, AP2 is present at the SAM, where it represses flowering, in part via direct repression of *SOC1*. Long days constitute an inductive signal that triggers *SOC1* expression at the SAM. SOC1 is a negative regulator of *AP2* expression, and reduces *AP2* expression during floral transition. However, before full repression of *AP2* it acts during floral transition to increase SAM size, height and width giving rise to a dome-shaped SAM. AP2 only acts at this transient stage to greatly alter SAM size and shape despite being expressed during the vegetative stage; therefore, we propose that other processes (here labelled as X) are induced during floral transition that are required for AP2 to cause morphological changes at the SAM. The full repression of *AP2* expression by SOC1 terminates the increase in SAM size and changes in shape associated with floral transition.

The effect of AP2 on SAM size and morphology is tightly temporally regulated. AP2 only detectably promotes changes in SAM size and shape during the early stages of floral transition, suggesting that the promotion of meristem size by AP2 requires other factors (represented as X in **Fig. 6**) that are induced during floral transition and whose early expression must also depend on SOC1. Moreover, even in *rAP2-V* plants, the SAM does not continue to increase in height and width after 17 LD, despite the continued expression of AP2 and SOC1, suggesting that the effect of AP2 on SAM size is actively terminated or is dependent on processes (e.g. X in Figure 6) that occur early during floral transition. AP2 may influence SAM development by increasing *WUS* transcription, because it does so in floral meristems towards the end of floral development^48^ and also in the SAM at the end of inflorescence development when shoot growth has terminated^37^. However, AP2 can promote inflorescence meristem size when expressed either in the *WUS* or *CLV3* domains, and *WUS* expression can be influenced by many meristem regulators^8,24–28,49^; thus, the mechanism by which AP2 increases *WUS* expression might be indirect and complex. Similarly, because AP2 is expressed during vegetative development but does not increase SAM size at that stage, it is unlikely to be sufficient to activate *WUS* expression. Moreover, SOC1, which is repressed by AP2, reduces the expression of enzymes involved in gibberellin biosynthesis and catabolism in the SAM, and this may alter the rate of meristem growth in *soc1* mutants^5^. The role of AP2 in increasing SAM size and altering its morphology during floral transition may therefore be complex and involve more than one regulatory mechanism.

A feature of our model is that the mutual repression of *AP2* and *SOC1* determines the timing of floral transition and changes in SAM size and shape. This type of direct reciprocal repressive motif has been characterized in developmental processes in animals^50–52^, and is proposed to sharpen and steepen spatial boundaries of gene expression^53^. AP2 and SOC1 are expressed in a similar spatial pattern throughout the SAM, but show different temporal patterns, with AP2 being expressed before SOC1. Thus, the mutual repression of SOC1 and AP2 determines the time interval during which both factors are expressed, which limits the stage during which cauline leaves are formed and AP2 promotes changes in SAM morphology.

## Materials and methods

### Plant material and growth conditions

All plants in this study were *Arabidopsis thaliana* Columbia-0 (Col-0) background. Mutant alleles were previously described: *ap2-12*^32^ and *soc1-2*^14^. The following transgenic lines were used: *AP2::AP2-VENUS #13*^40^*, AP2::rAP2VENUS #A6*^8^ and *SOC1::SOC1-GFP*^44^. The *ap2-12 soc1-2* double mutant (previously published in ^32^ and reconstructed here) and the *AP2::AP2-VENUS #13*/*soc1-2 ap2-12* and *SOC1::SOC1-GFP/soc1-2 ap2-12* genotypes were generated in this study via crossing. Plants were grown on soil under controlled conditions of SDs (8 h light/16 h dark) and LDs (16 h light/8 h dark).

### Phosphinotricin (PPT) resistance assay

The identification of PPT-resistant plants was performed on agar plates using Murashige & Skoog (MS) medium containing PPT similar to^40^, but using 1× MS medium containing 15 mg mL^-1^ PPT (without sucrose), and placing the plates with leaves in continuous light at 21°C for at least 5 days.

### Confocal imaging

Shoot apices at different developmental stages were dissected under a stereomicroscope and fixed with 4% (v/v) paraformaldehyde (PFA; Electron Microscopy Sciences). The fixed samples were washed twice for 1 min in phosphate-buffered saline (PBS) and cleared with ClearSee^54^ for 3–4 days at room temperature. Before imaging lines containing a fluorescent reporter, samples were kept in PFA for 2 h at room temperature after fixation and were then transferred to PBS for 2 days and then to ClearSee for 3–4 days. The cell wall was stained with Renaissance 2200 [0.1% (v/v) in ClearSee]^55^ for at least 1 day.

Confocal microscopy was performed with a TSC SP8 confocal microscope (Leica) for cell segmentation and SAM morphology quantification, where Renaissance was excited at 405 nm and image collection was performed at 435–470 nm (**Fig. 1**, **Fig. 3**a, **Fig. 4**e–f, Supplementary Figure 1, Supplementary Figure 2, Supplementary Figure 4, Supplementary Figure 5, Supplementary Figure 7 and Supplementary Figure 8). The spatial patterns of AP2:VENUS and rAP2:VENUS were examined using a LSM780 confocal laser scanning microscope (Zeiss), where VENUS was excited at 514 nm and the signal was detected at 517–570 nm (**Fig. 2**, Supplementary Figure 3). The protein patterns of AP2:VENUS and SOC1:GFP under continuous LDs at the SAM were acquired with a Stellaris 5 confocal microscope (Leica) for fluorescence quantification (**Fig. 3**d–e, **Fig. 5**, Supplementary Figure 6, Supplementary Figure 11). VENUS and GFP were excited at 515 nm and 488 nm, and the signal was detected at 520–600 nm and 500–557 nm, respectively. For all time courses where protein accumulation patterns were determined, Renaissance signal was detected using similar parameters as mentioned earlier for the segmentation analyses.

### Cell segmentation and SAM morphology quantification

The z-stacks of SAMs were acquired with a step size of 0.4 µm and were converted to TIF files with Fiji. MorphoGraphX (MGX) software (https://morphographx.org/)^56,57^ was used to extract the surface of the meristem and to project the Renaissance signal of the cell wall from the outer cell layer (i.e. L1), which was used to segment the images. Cells were segmented using the “auto-segmentation” function and corrected manually. The geometry of the surface was displayed as Gaussian curvatures with a neighbouring radius of 10 μm. The boundary between the meristem and the developing primordia was defined by a negative Gaussian curvature, then the area of each of the cells in the SAM was extracted. The meristem area was calculated as the sum of the areas of the cells that comprised the meristem.

To quantify the morphology of the meristem, its height and width were estimated (Supplementary Figure 2a). For this, the orthogonal views from the z-stacks were generated and were used to estimate the meristem height and the width according to the following criteria: (1) the height aligned with the apical–basal axis, (2) the width was perpendicular to the height and (3) the width was measured from the most apically visible primordium. The measurements performed on each orthogonal view were considered as technical replicates; thus, the plotted values corresponded to the means of the two estimations of each of the measured parameters. The parabolas to represent meristem morphology were fitted in an XY-coordinate system (Supplementary Figure 2c) using the formula in Supplementary Figure 2d. For representation purposes, the parabolas were coloured according to the identity of primordia that were formed at the SAM periphery (**Fig. 1**d, **Fig. 3**a, **Fig. 4**f, Supplementary Figure 2h–j).

### Fluorescence quantification

Confocal fluorescence z-stacks were processed and analysed using a Matlab custom code (https://gitlab.com/slcu/teamHJ/pau/RegionsAnalysis). The main goal of this analysis was to extract reproducible measures of fluorescence intensity within the SAM. A normalised fluorescence intensity measure was computed as a proxy for the concentration of protein at the meristem upper region. To do that, a semi-automatic pipeline was developed, which is described as follows.

Due to the difference in the resolution between the xy plane and the *z*-direction (depth), the z-stack was resized by increasing the number of slices in the z direction through bicubic interpolation to obtain a homogeneous volumetric resolution.

To exclude fluorescence signals outside the region of interest and quantify only the fluorescence intensity within the meristematic region, a pre-processing step was performed: a 3D paraboloid mask was constructed using the curvature of the meristem (Supplementary Figure 11). First, a stack-slice interval that contained the apex of the meristem was selected and the cell wall signal present within this interval was projected in each orthogonal plane (xy and yz) (Supplementary Figure 11a). Then, two curved lines following the parabolic outline of the SAM were drawn in the xy and yz planes, in the sum of slice projection of each plane (Supplementary Figure 11b). Later, a parabolic fitting of the two drawn lines was performed (Supplementary Figure 11c), which considered tilting of the drawn curve lines. From the two orthogonal parabolas fitted for each z-stack, the apex was computed to derive the equation for the 3D paraboloid. The *z_o_* coordinate of the paraboloid was determined from averaging the apices of the orthogonal parabolas (Supplementary Figure 11d). The parameter *a* in the parabola equation, i.e the curvature, was used to substitute the denominator terms in the paraboloid equation (*c*_1_ ^2^ and *c*_2_ ^2^; Supplementary Figure 11d) so that the paraboloid equation matched the linear and quadratic terms of each of the equations at *y*=*y*_o_ and *x*=*x*_o_, respectively (Supplementary Figure 11d). Because the z-stack did not always include the beginning and end of the meristem in the yz plane (lateral view), only the xy curvature was used (*c* ^2^ = *c* ^2^). Finally, to extract the fluorescence signal within the SAM using the 3D paraboloid, a 2D parabolic mask was created for each of the z-stack slices, and, in each slice of the z-stack, all intensity values of the pixels outside the paraboloid were set to 0 (Supplementary Figure 11d– e).

To exclude any fluorescence signal at the boundaries of the SAM and primordia, the paraboloid curvature was increased with respect to the original (Supplementary Figure 11d) such as *a*’=a/α, being *a*’ the curvature of the new paraboloid and α*<*1 μm the image resolution (Supplementary Figure 11d).

A concentration of fluorescence intensity measure was computed as the fraction between the total intensity (sum of the voxels’ intensity) and the total volume (sum of the voxels’ volume) in the upper region of the paraboloid with increased curvature. This upper paraboloid region was delimited between the 3D paraboloid itself and a transversal plane set at a distance of 20 μm or 50 μm from the paraboloid apex (Supplementary Figure 11e). For *SOC1::SOC1-GFP/soc1-2 ap2-12* fluorescence quantification a Gaussian filter (sigma = 2.5) was applied in the region within the paraboloid. For representation purposes, the values for the concentration of fluorescence intensity were divided by 1000.

### Gene expression and whole-transcriptomic RNA-sequencing analysis

Shoot apices of Col-0 and *ap2* mutants were dissected under a stereo microscope at 10, 12, 14 and 17 days in LD conditions in three independents biological replicates. Total RNA was extracted using the RNeasy Plant Mini Kit (Qiagen, USA) and subjected to DNase treatment using the TURBO DNase (Invitrogen). Poly(A) RNA enrichment, library preparation, and sequencing were carried out at the MPIPZ Genome Center, Cologne, Germany using the following conditions: The RNAs were processed by poly-A enrichment followed by application of basic components of “NEBNext Ultra II Directional RNA Library Prep Kit for Illumina” with a homebrew barcoding regime. Sequencing was performed on a HiSeq3000 sequencer by sequencing-by-synthesis with 1× 150 bp single-read length. Sequence reads were preprocessed to remove any residual adaptors with CutAdapt, and the low-quality bases (Q<15) were trimmed from the ends with Trimmomatic^58,59^. Only reads with a minimum length of 50 nucleotides were kept. Salmon was used to quantify the abundance of transcripts from the Arabidopsis reference genome Reference Transcript Dataset for Arabidopsis (including guanine/cytosine bias, unstranded samples)^60,61^. Fragments Per Kilobase of transcript per Million (FPKM) values and corrected *p*-values were obtained using DESeq2 by comparing Col-0 to *ap2* in each time point using standard settings. Differentially expressed genes (DEGs) were defined in each comparison via DESeq2^62^ (i.e. adjusted *p*-value < 0.05 and absolute Log_2_ Fold Change > 1).

## Acknowledgements

We thank C. Ferrándiz and the members of the laboratory of G.C. for engaging scientific discussions. We thank J. Chandler for critical reading of the article. We are grateful to B. Huettel and the staff from the Max Planck Genome Centre for their support during the RNA-sequencing experiment. We thank the staff from the cultivation facilities of the MPIPZ. M.C. received a post-doctoral fellowship from the Alexander von Humboldt foundation and K.W. received a studentship from the China Scholarship Council. GC receives funding from the Cluster of Excellence CEPLAS (EXC 2048/1 Project ID: 390686111) and a Core Grant from the Max Planck Society.

## Author contributions

E.B.GdO. and G.C. conceived and designed the study. A.V., E.B.GdO. and E.S. produced the RNA-Seq data. E.B.GdO, G.R-M., M.C and Y.L.S. performed confocal imaging and the subsequent analyses. K.W. introduced the AP2 translational reporter in the *ap2-12 soc1-2* mutant background. Y.L.S. and S.S. provided technical support. A.V., P.F-J. and G.C. provided scientific guidance throughout the project. E.B.GdO and G.C. wrote the manuscript. All the authors discussed the results and commented on the manuscript.

## Materials and correspondence

Correspondence to George Coupland.

## Supplementary Figure legends

Supplementary Figure 1. Supplementary data for Fig. 1. (a) Quantification of the changes in cell area in the meristem region of Col-0, *ap2-12* and *rAP2-VENUS* SAMs during the time course in Fig. 1. White dots represent the median value for each genotype. Significant differences among genotypes were determined for each time point via one-way ANOVA, followed by Tukey post-hoc comparisons (*p* < 0.05). Data sets that share a common letter do not differ significantly. The colour of the letters and the shapes denoting the data (i.e. violin plots) correspond to the studied genotypes. *N* = 3–4 SAMs. (b–d) Top view images from segmented meristems of (b) Col-0, (c) *ap2-12* and (d) *rAP2-V* at +9 LD. Asterisks indicate representative morphologies of developing organs: leaves (black), cauline leaves subtending axillary branches (yellow) and flowers (white). Scale bar = 50 μm.

Supplementary Figure 2. Method to quantify changes in SAM morphology by measuring height and width of the shoot meristem. (a) Lateral view of the heatmap quantification of cell area in the meristem region of Col-0, *ap2-12* and *rAP2-VENUS* SAMs from 2-week-old plants grown under short day (SD) conditions (2 wSD) and then transferred to continuous long days (LDs). White asterisks indicate the first time point in which floral primordia were detected in the analysis of the corresponding genotype. Scale bar = 50 μm. (b) Two orthogonal views were generated from each z-stack. In each orthogonal view, height and width were measured following the specified criteria in materials and methods. The two measurements for each parameter were considered as technical replicates; therefore, the mean of these values was used for further analysis. (c) Normalised height and width of Col-0 to the first time point of the time course shown in Supplementary Figure 2f–g. The colour of the dots corresponds to the represented parameter. (d–e) Adjustment of the measured height and width to a parabola. (d) Representation of height and width in a cartesian coordinate system. Notice that three points can be placed at the extremes of these measurements, which define a unique parabola. (e) General equation to represent a parabola from measured height and width. (f–g) Measurement of (b) width and (c) height of the SAM in 2wSD-grown plants that were then transferred to LDs. The horizontal bars represent the median value for each genotype. Significant differences intra-time point among genotypes were determined via one-way ANOVA, followed by Tukey post-hoc comparisons (*p* < 0.05). Data sets that share a common letter do not differ significantly. The colour of the dots and the letters correspond to the genotype. (h–j) Col-0 meristems producing (h) rosette leaves (10 LD), (i) cauline leaves (14 LD) and (j) flowers (17 LD) at their flanks. Arrowheads mark the characteristic morphology of the developing organs that are indicated in the description of each image. Scale bar = 50 μm.

Supplementary Figure 3. Images without clipping depicting pattern of protein accumulation of *AP2::AP2:VENUS* in *ap2-12* and *AP2::rAP2:VENUS* in Col-0 at the SAM of 2-week-old plants grown under short day (SD) conditions (2 wSD) and then transferred to continuous long days (LDs) shown in Fig. 2. The shape of the acquired meristem and its peripheral organs is indicated with a dotted white line. Scale bar = 50 μm.

Supplementary Figure 4. Measured (a) height and (b) width of the Col-0, *ap2-12* and *rAP2-VENUS* meristems represented in Fig. 3a. The horizontal bars represent the median value for each genotype. Significant differences intra-time point among genotypes were determined via one-way ANOVA, followed by Tukey post-hoc comparisons (*p* < 0.05). Data sets that share a common letter do not differ significantly. The colour of the dots and the letters correspond to the genotype. *N* = 10–21.

Supplementary Figure 5. Segmentation analysis of the SAM of Col-0, *ap2-12* and *rAP2-VENUS* under continuous long days (LDs). (a) Top view of the heatmap quantification of cell area in the meristem region. White asterisks indicate the first time point in which floral primordia were detected in the analysis of the corresponding genotype. Scale bar = 50 μm. *N* = 4. (b–d) Quantification of (b) meristem area and (c) cell number and (d) cell area in the meristem region. The (b–c) horizontal bars and the (d) white dots represent the median value for each genotype. Significant differences intra-time point among genotypes were determined via one-way ANOVA, followed by Tukey post-hoc comparisons (*p* < 0.05). Data sets that share a common letter do not differ significantly. The colour of the dots, the letters and the shapes denoting the data (i.e. violin plots) correspond to the genotype.

Supplementary Figure 6. Images without clipping depicting the pattern of protein accumulation of *SOC1::SOC1:GFP* in *soc1-2* and *soc1-2 ap2-12* mutant backgrounds at the SAM of plants grown under continuous long day (LD) conditions, shown in Fig. 3d. The shape of the acquired meristem and its peripheral organs is indicated with a dotted white line. Scale bar = 50 μm. (b) Quantification of SOC1:GFP concentration of fluorescence intensity (total fluorescence divided by volume) at the shoot apex (from the tip to 20 μm deep in the basal direction) in *soc1-2* and *soc1-2 ap2-12* mutant backgrounds during continuous LDs. The dots are coloured according to the mutant background of the analysed plant. Comparisons intra-time point between genotypes were performed via Wilcoxon rank sum test. Statistical significance (*p* < 0.075) is indicated with an asterisk. *N* = 6–10.

Supplementary Figure 7. Measured (a) height and (b) width of the meristems of Col-0, *ap2-12*, *soc1-2* and *soc1-2 ap2-12* represented in Fig. 4e. The horizontal bars represent the median value for each genotype. Significant differences intra-time point among genotypes were determined via one-way ANOVA, followed by Tukey post-hoc comparisons (*p* < 0.05). Data sets that share a common letter do not differ significantly. The colour of the dots and the letters correspond to the genotype.

Supplementary Figure 8. Quantification of (a) meristem area, (b) cell number and (c) cell area in the meristematic region shown in Fig. 4f. The horizontal bars (a–b) and white dots (c) represent the median value for each genotype. Significant differences intra-time point among genotypes were determined via one-way ANOVA, followed by Tukey post-hoc comparisons (*p* < 0.05). Data sets that share a common letter do not differ significantly. The colour of the dots, the shapes denoting the data (i.e. violin plots) and the letters correspond to the genotype. *N* = 4.

Supplementary Figure 9. GBrowse traces of mapped ChIP-Seq reads for AP2 and SOC1. Previously published ChIP-seq data for AP2 ^32^ and SOC1 ^44^ were reanalyzedBinding of (a) AP2 to the promoter region of *SOC1* and (b) SOC1 to the promoter region of *AP2*. Gene models are shown under the upper scale bar. The blue horizontal bars below the track of the gene models delimit the position of the peak regions. The coverage of the mapped reads is represented in the lower tracks in each subfigure by grey bars.

Supplementary Figure 10. Supplementary data for Fig. 5. (a) Images without clipping depicting the pattern of protein accumulation of *AP2::AP2:VENUS* in *ap2-12* and *ap2-12 soc1-2* mutants at the SAM of plants grown under continuous long day (LD) conditions shown in Fig. 5. The shape of the acquired meristem and its peripheral organs is indicated with a dotted white line. Scale bar = 50 μm. (b) Quantification of AP2:VENUS concentration of fluorescence intensity (total fluorescence divided by volume) at the shoot apex (from the tip to 20 μm deep in the basal direction) in *ap2-12* and *soc1-2 ap2-12* mutants during continuous LDs. The dots are coloured according to the mutant background of the analysed plant. The horizontal bars represent the median value for each genotype. Comparisons intra-time point between genotypes were performed via Wilcoxon rank sum test. Statistical significance (*p* < 0.075) is indicated with an asterisk. *N* = 5–10.

Supplementary Figure 11. Fluorescence quantification explanation pipeline. (a) SAM fluorescence confocal microscopy images were obtained from the lateral side. In the image: slices from z-stack meristem from a 12-day-old plant grown under LDs containing the *AP2::AP2:VENUS* reporter. The membrane marker channel is displayed in white and the AP2-VENUS fluorescence signal is coloured according to the Fire colour look-up table in Fiji. Scale bar = 20 μm. The acquisition parameters are described in Materials and Methods. (b) A curved line (in red) marks the meristem outline until the axils of the primordia on the sum of slice projections of certain z-stack slice intervals. (c) Parabolic fitting of the previous parabola. Triangles mark the beginning and end points of the drawn curve line (white) and fitted parabola (black). Circles represent the apex position for the drawn curve line (white) and fitted parabola (black). (d) 3D paraboloids built from the previously extracted orthogonal parabolas. In orange, paraboloid using the original curvature values, with the extracted orthogonal parabolas outlined on top of it; in purple, paraboloid with higher curvature to focus on the central area of the meristem along its longitudinal axis. The purple paraboloid is used to generate a mask such that when multiplied to a given z-stack, the intensity values of pixels whose positions lie outside the paraboloid are set to zero. P.p. = pixel position. (e) Selected 3D-paraboloid section of interest (0 - 50μm distance from the apex coordinates) in which the fluorescence intensity is quantified. f) Main set of equations describing the 2D parabola in the xy-plane, its maximum position and the 3D paraboloid. The 2D equation of the parabola in the zy plane is omitted for simplicity.

## Supplementary table legends

Supplementary Table 1. List of differentially expressed genes in *ap2-12* vs. Col-0 at least at one time point during the RNA-seq time course.

Supplementary Table 2. List of differentially expressed genes in *ap2-12* vs. Col-0 at 14 LD.

Supplementary Table 3. List of differentially expressed genes in Col-0 at 14 LD vs. 10 LD.

Supplementary Table 4. List of genes that are present in Supplementary Table 2 and Supplementary Table 3.

Supplementary Table 5. List of direct direct target genes of AP2 identified by ChIP-Seq analysis in^32^.

