## Supplementary Figures for "A regulatory gene network that couples floral transition to shoot apical meristem morphology in Arabidopsis"

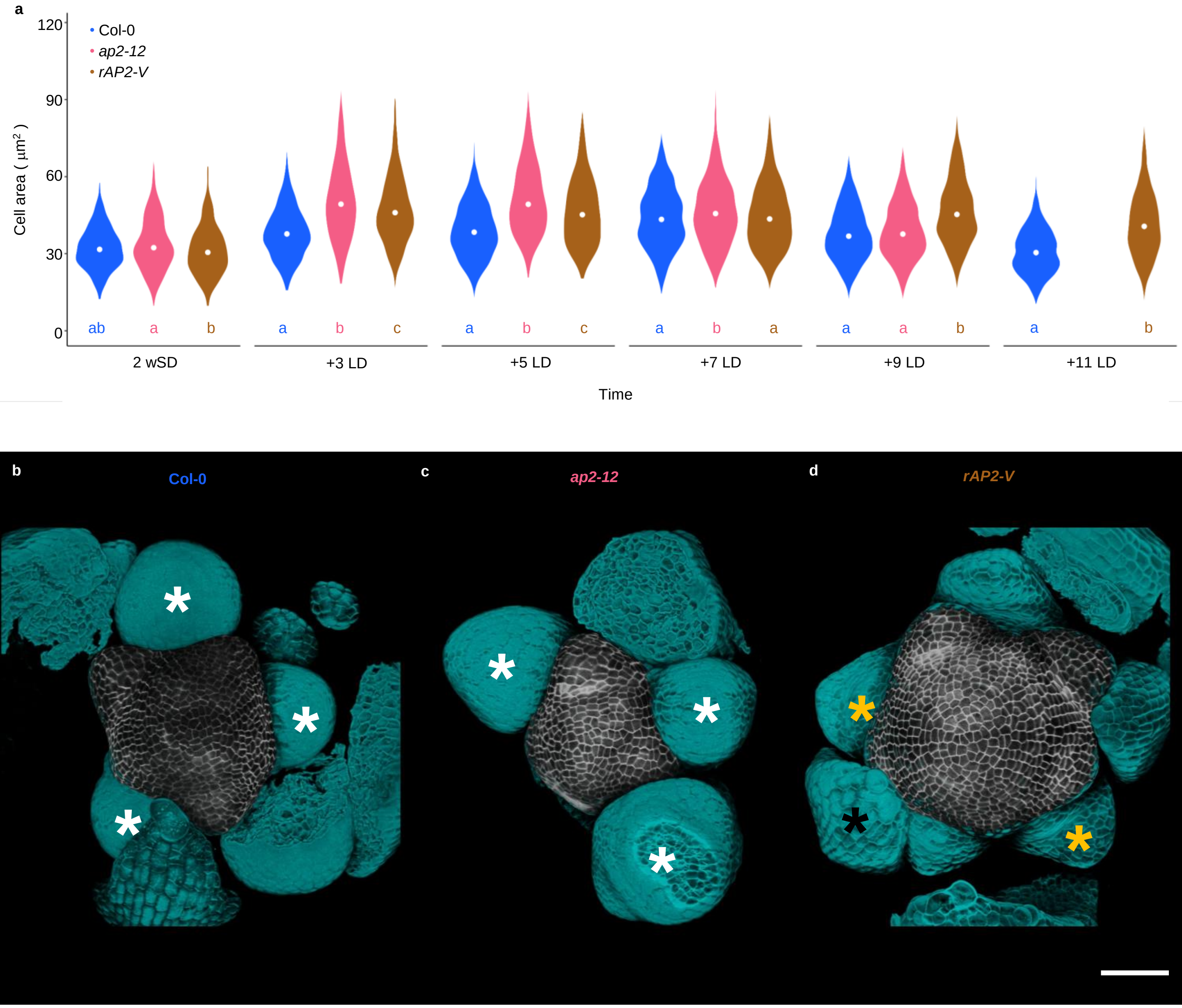


**Supplementary Figure 1**

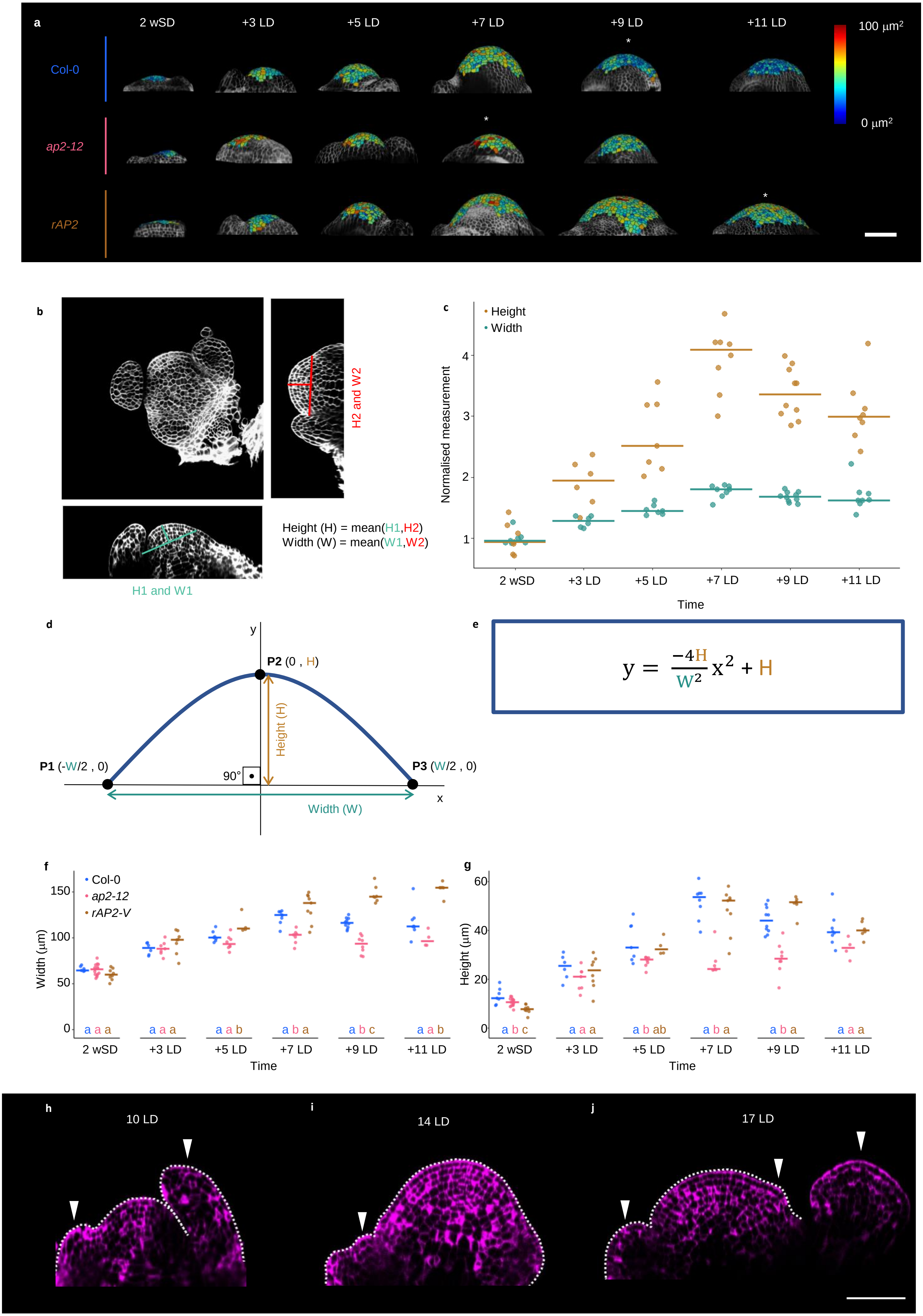


**Supplementary Figure 2**

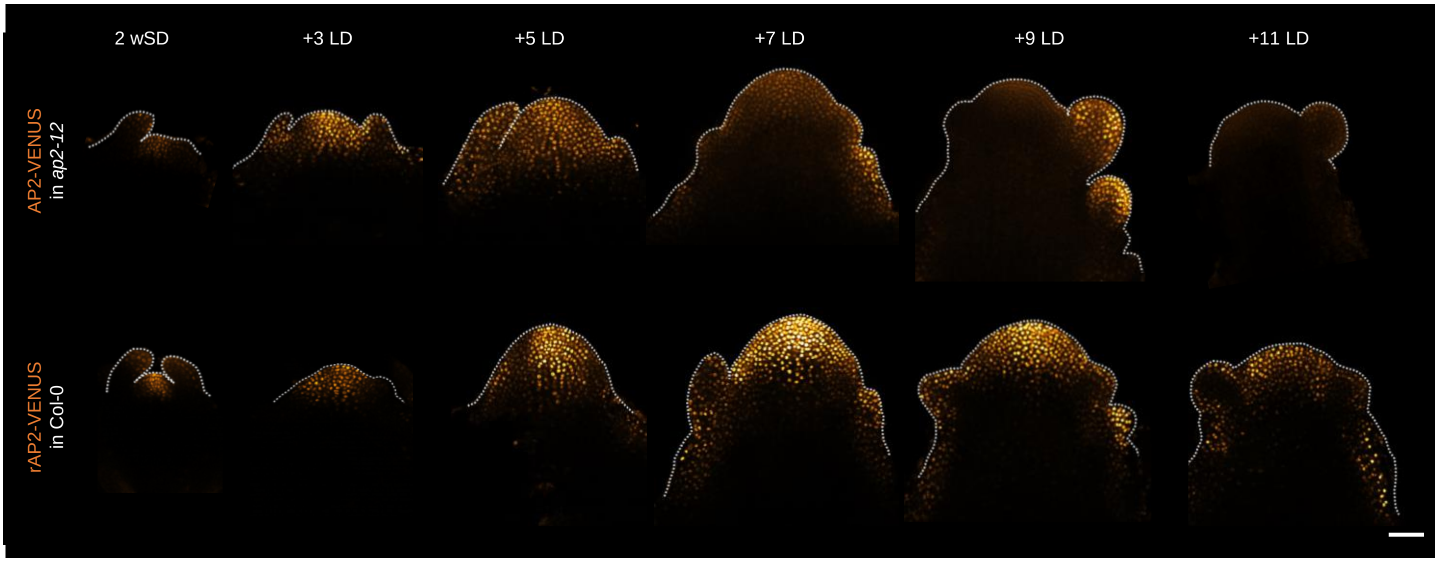


**Supplementary Figure 3**

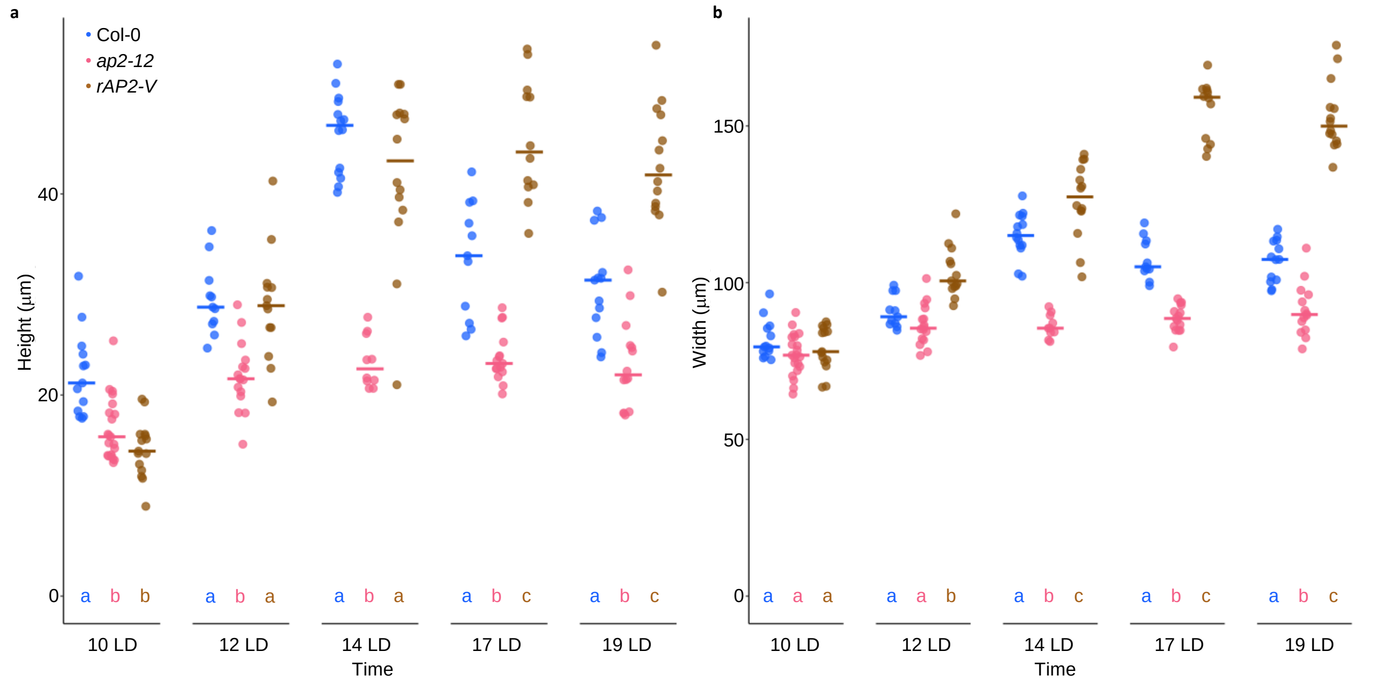


**Supplementary Figure 4**

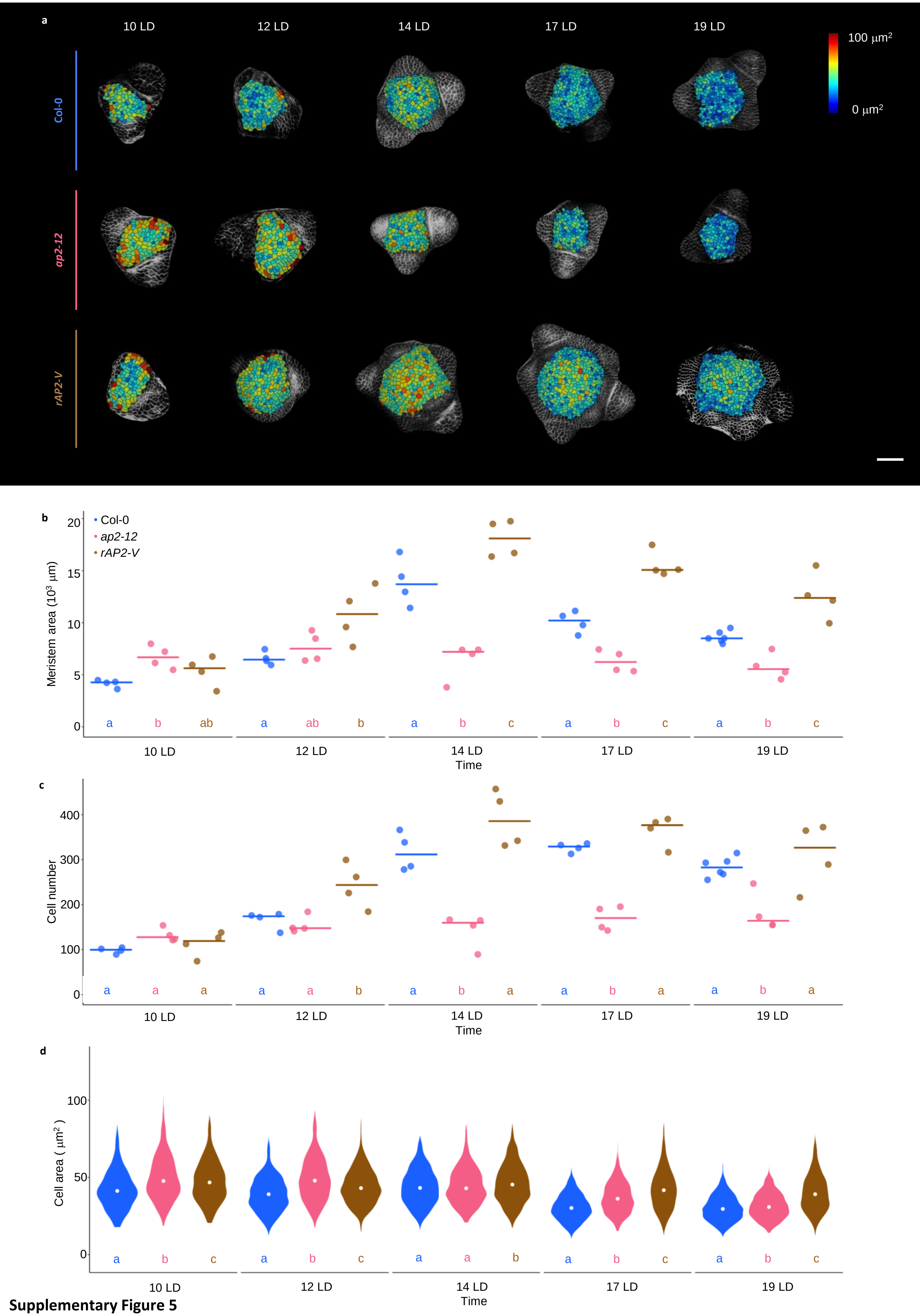


**Supplementary Figure 5**

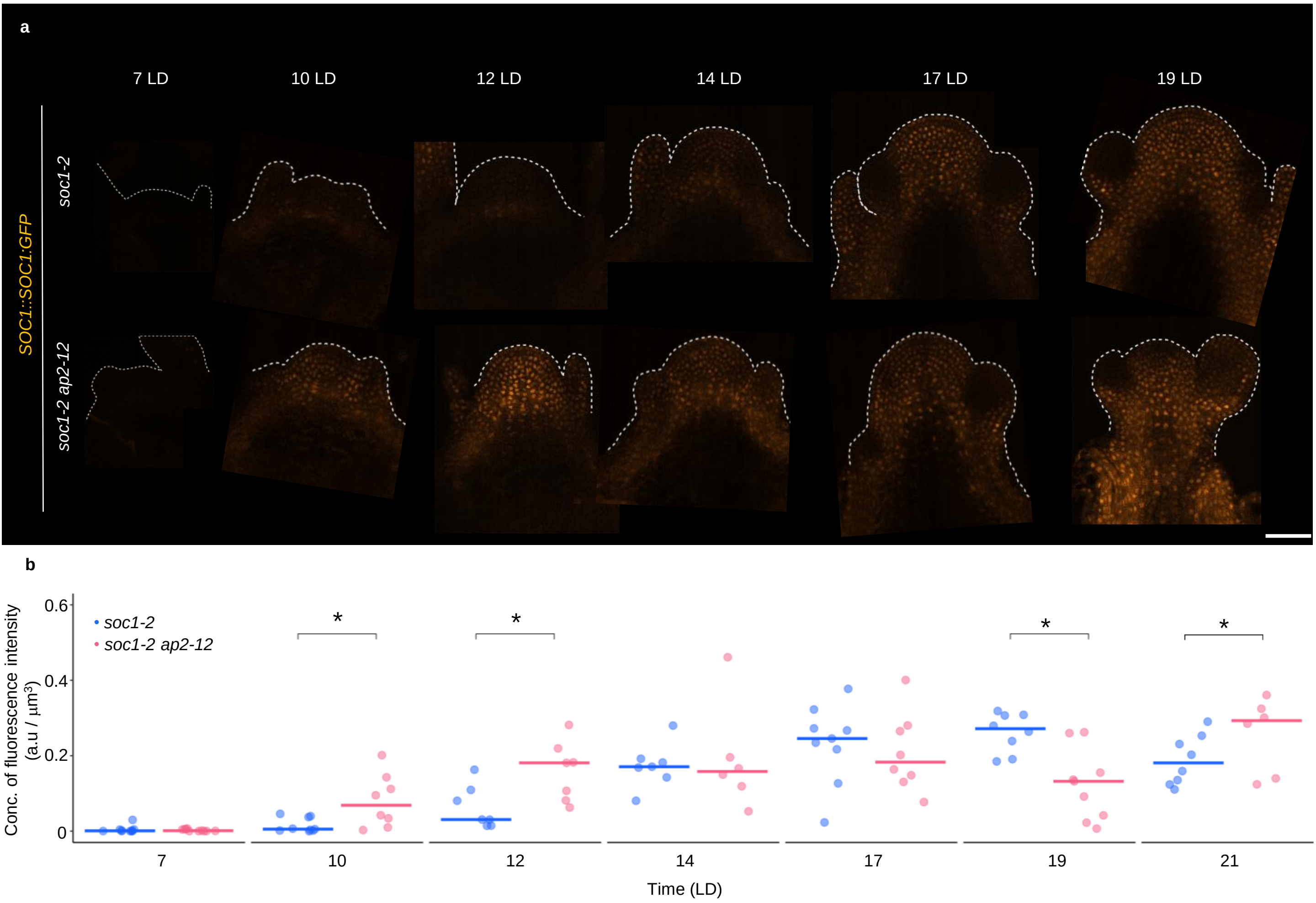


**Supplementary Figure 6**

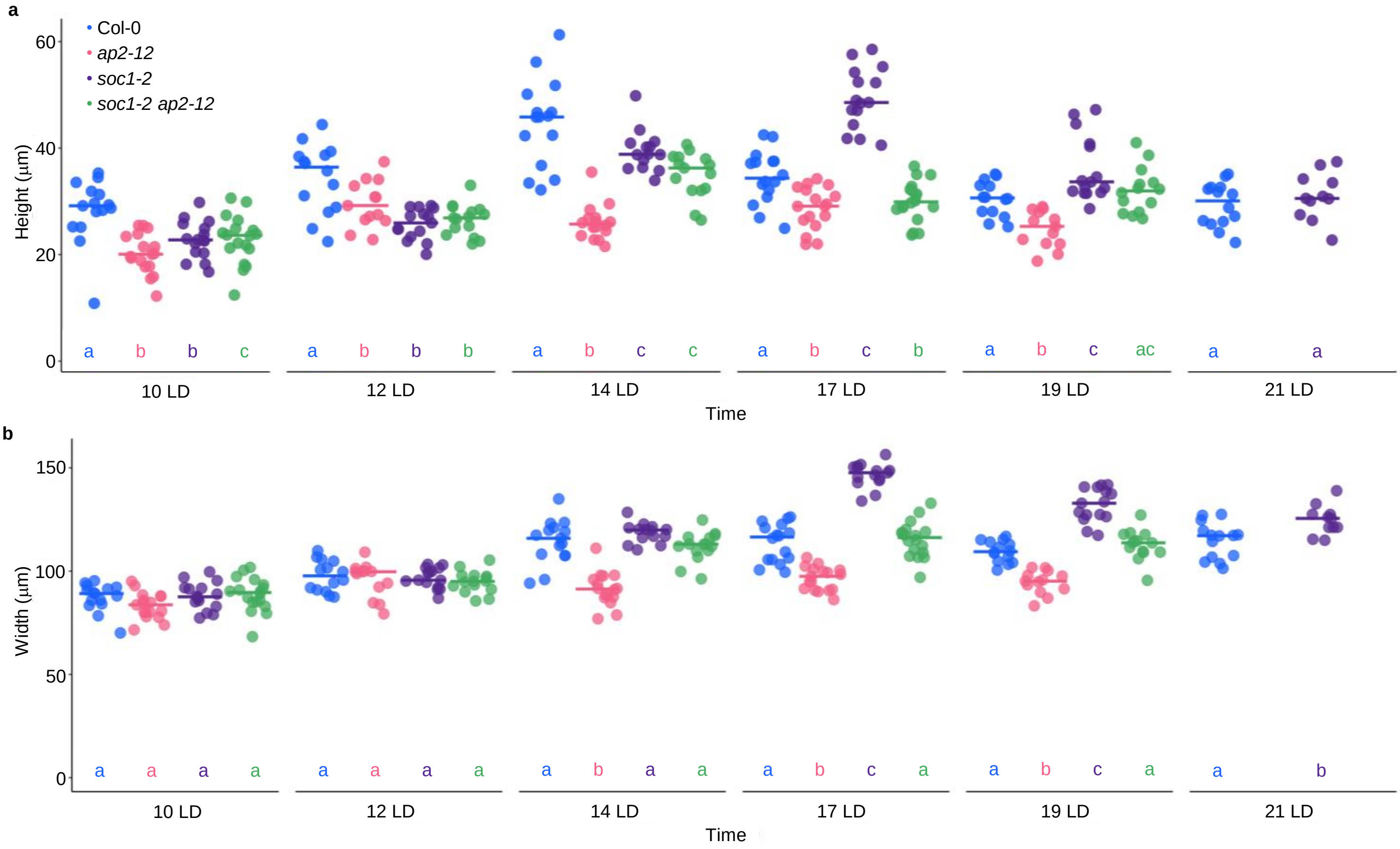


**Supplementary Figure 7**

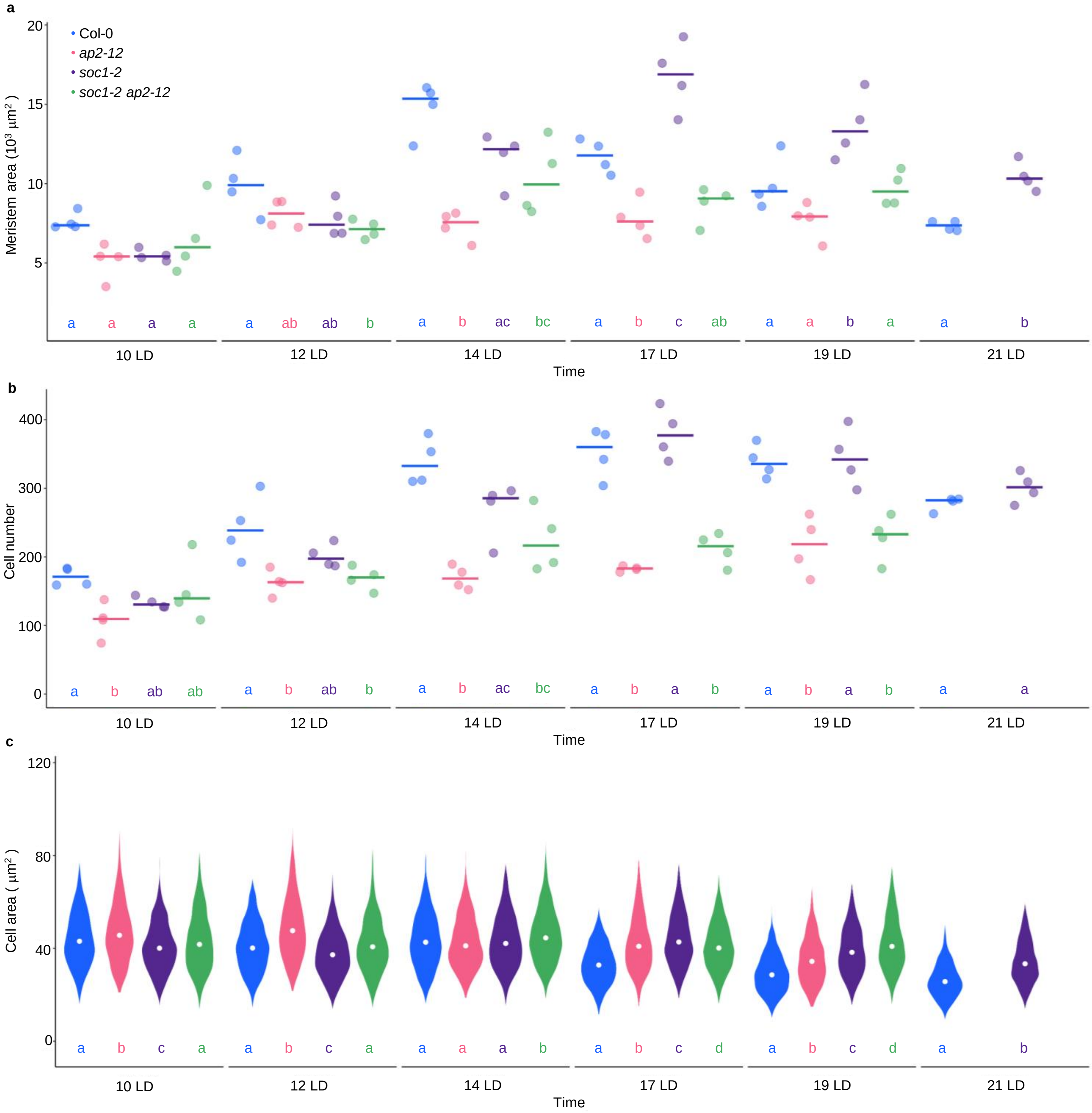


**Supplementary Figure 8**

a

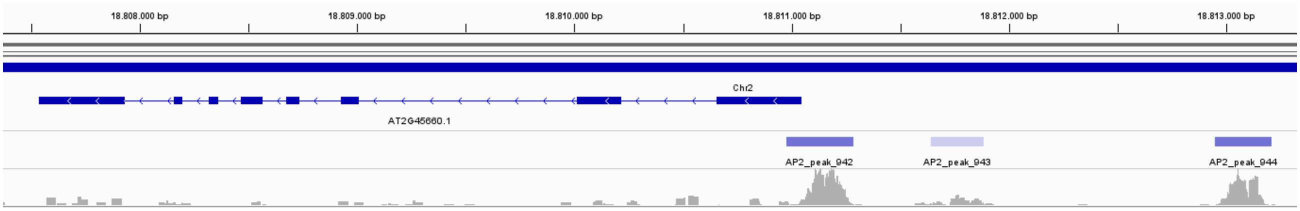


b

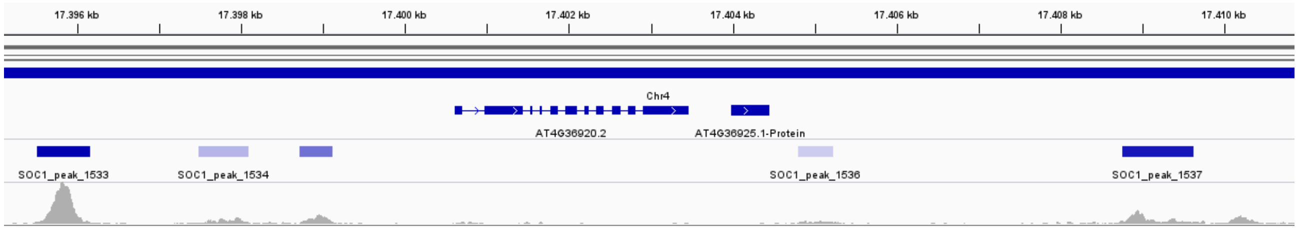


**Supplementary Figure 9**

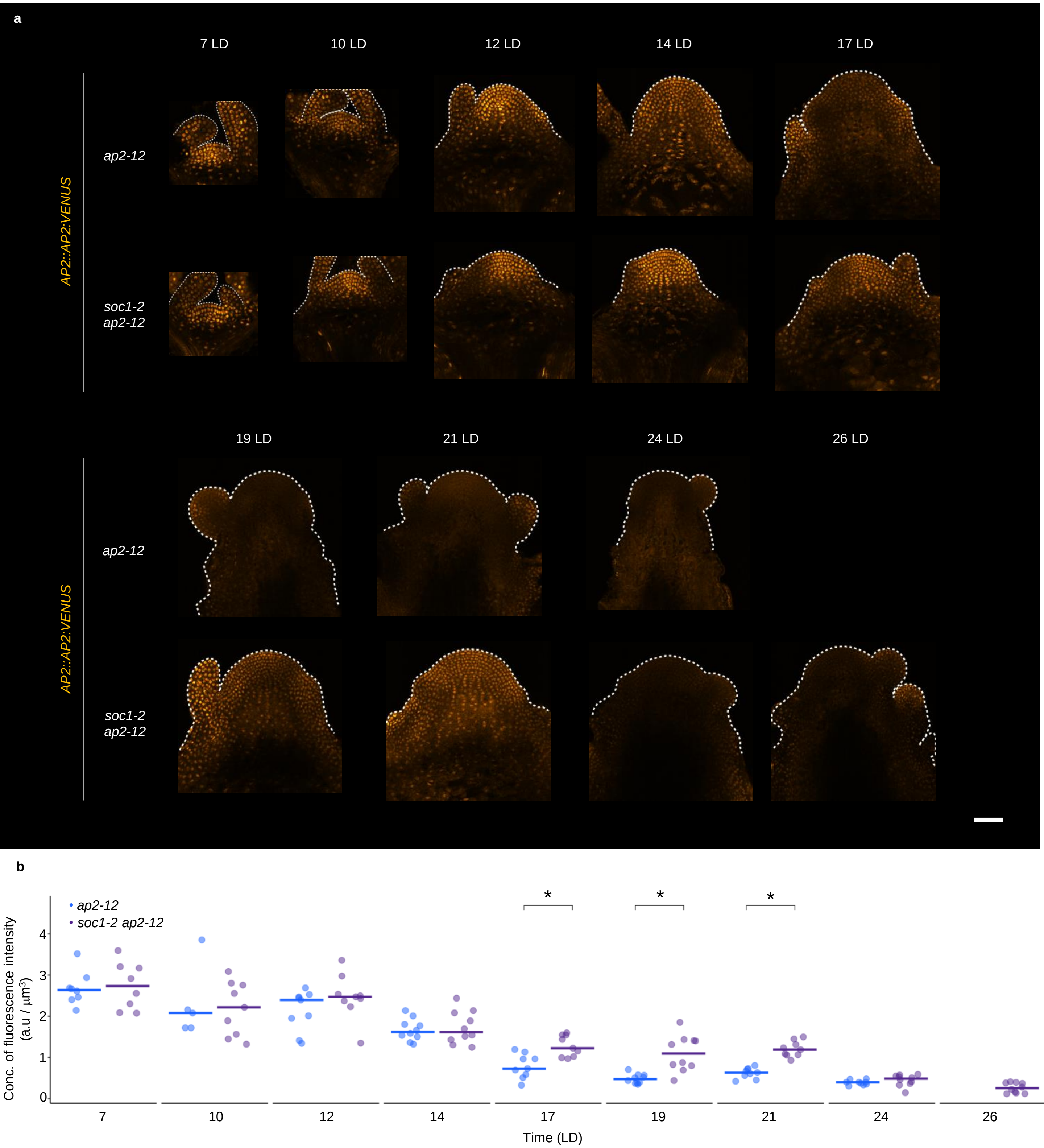


**Supplementary Figure 10**

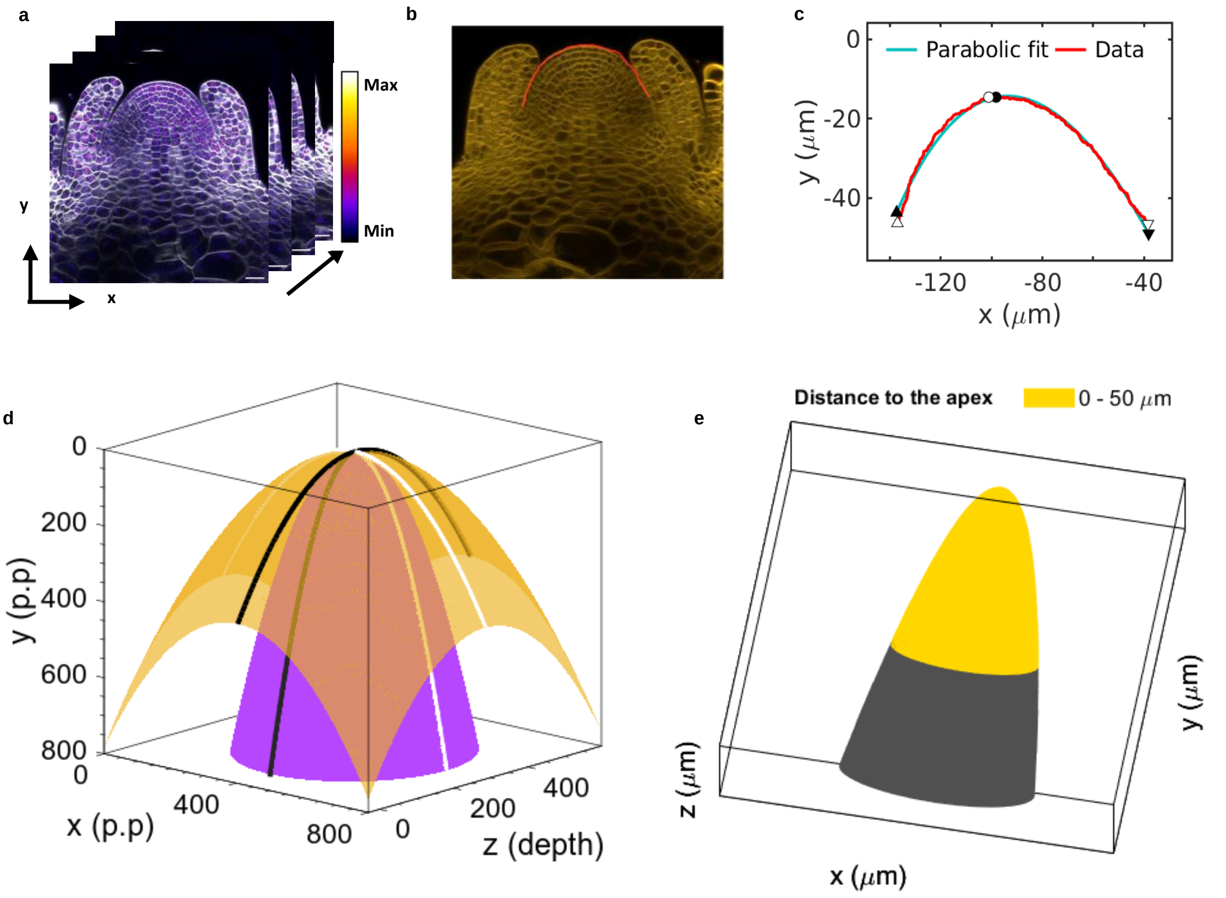


**Parabola, pos. max. & paraboloid**

$$y = a(x - x_o)^2 + b(x - x_o) + c \qquad x_{max} = x_o - \frac{b}{2a} \qquad z - z_o = -\frac{(x - x_o)^2}{c_1^2} - \frac{(y - y_o)^2}{c_2^2}$$

**Supplementary Figure 11**
